# Mapping the blood vasculature in an intact human kidney using hierarchical phase-contrast tomography

**DOI:** 10.1101/2023.03.28.534566

**Authors:** Shahrokh Rahmani, Daniyal J Jafree, Peter D Lee, Paul Tafforeau, Joseph Brunet, Sonal Nandanwar, Joseph Jacob, Alexandre Bellier, Maximilian Ackermann, Danny D Jonigk, Rebecca J Shipley, David A Long, Claire L Walsh

## Abstract

The architecture of the kidney vasculature is essential for its function. Although structural profiling of the intact rodent kidney vasculature has been performed, it is challenging to map vascular architecture of larger human organs. We hypothesised that hierarchical phase-contrast tomography (HiP-CT) would enable quantitative analysis of the entire human kidney vasculature. Combining label-free HiP-CT imaging of an intact kidney from a 63-year-old male with topology network analysis, we quantitated vasculature architecture in the human kidney down to the scale of arterioles. Although human and rat kidney vascular topologies are comparable, vascular radius decreases at a significantly faster rate in humans as vessels branch from artery towards the cortex. At branching points of large vessels, radii are theoretically optimised to minimise flow resistance, an observation not found for smaller arterioles. Structural differences in the vasculature were found in different spatial zones of the kidney reflecting their unique functional roles. Overall, this represents the first time the entire arterial vasculature of a human kidney has been mapped providing essential inputs for computational models of kidney vascular flow and synthetic vascular architectures, with implications for understanding how the structure of individual blood vessels collectively scales to facilitate organ function.

## INTRODUCTION

The vascular system of the kidney is highly specialised, serving multiple functions including delivery of oxygen and nutrients to the organ’s parenchyma whilst also facilitating plasma ultrafiltration and solute reabsorption. Despite only comprising approximately 1% of body weight, the kidney receives up to 20% of cardiac output [1]. Blood enters the kidney through the kidney artery which branches from the abdominal aorta and enters the kidney hilum. Once within the kidney, the kidney arteries divide hierarchically, first into segmental arteries which pass through the kidney pelvis, then branching into interlobar branches which pass through columns between the pyramids of the kidney medulla. At the distal end of the kidney columns, interlobular arteries branch into arcuate arteries that arch around the outer surface of the kidney pyramids. From these, the interlobular vessels branch and penetrate into the surrounding kidney cortex before finally terminating at efferent arterioles [1]. This complex network perfuses specialised glomerular capillaries for plasma ultrafiltration, before peritubular capillaries located in the cortex, and vasa recta located in the medullary pyramids facilitate dynamic solute exchange. Thereafter, venous return follows the arterial supply out of the organ [2].

Structural and molecular changes to the kidney vasculature are a common feature of kidney pathologies including multiple aetiologies of chronic kidney disease (CKD) and transplant rejection in both animal models and patients [3] Therefore, studying kidney vascular patterning has implications for understanding the basis of kidney function in health and disease, and also aids surgical planning for tumour resection, nephrectomy and transplantation. Vascular geometries also have a central role to play in computational models which underpin the creation of digital twins, such as through the creation of synthetic data [4], and flow modelling [4], [5], [6], [7], [8], which are playing an increasing role in biomedical research.

Vascular imaging of the kidney has advanced following technological innovations in micro-computed tomography (μCT) [9], [10], [11], magnetic resonance imaging (MRI) [12], ultrasound [13], lightsheet microscopy [14], [15] and photoacoustic imaging [16], [17]. These techniques have been used to generate quantitative analyses of vascular network geometry in intact kidneys of model organisms, particularly rodents where kidney diameter reaches up to 12 mm [18]. Comparatively human kidneys, with a diameter of approximately 5 cm [19] are far more challenging to image at high resolution whilst still intact. Corrosion casting of human kidneys has highlighted vascular heterogeneity and generated intricate 3D casts (down to 100 μm) but has provided limited quantitative or accessible digitized geometries of the vascular network [20]. Optical clearing and lightsheet microscopy have been used to quantify portions of the human kidney vascular network [21] however, it has not yet been possible to capture the intact vascular network of the human kidney without physical sectioning the tissue, beyond approximately six vessel divisions [22]. MRI has been used to quantify larger vessels both *in vivo* and post mortem [23], [24], but lacks the resolution capable of imaging small vessels and arterioles [23].

Due to these limitations, human kidney vascular network analysis is often predominantly focused on the very large, first three branches of the arterial tree [23], or on only small portions of the network [25]. Where multiscale modelling has been performed, parameters from rodent kidneys are assumed to be representative of human kidney vascular networks [4], [5], [8]. The semi-quantitative studies of human kidney vascular casts have shown large anatomical variation in even the segmental artery (first or second branch after the kidney artery) patterns [22], while smaller vessels such as arcuate arteries, interlobular arteries and afferent or efferent arterioles have not been assessed quantitatively at the organ scale.

Here, we show how a recently developed X-ray based non-destructive imaging technique - hierarchical phase-contrast tomography (HiP-CT), can be used to map and quantify the arterial vascular network of an intact human kidney down to the arteriolar level for the first time. HiP-CT is a technique which leverages the European Synchrotron Radiation Facility’s (ESRF) Extremely Brilliant Source (EBS); a high-energy fourth generation synchrotron source, to image intact human organs at unprecedented scale and resolution. Previously we have demonstrated the feasibility of applying HiP-CT to profile the human glomerular morphology and number across cubic centimetres of intact human kidney [26]. We now extend the use of this technology to extract and quantify the arterial network of an intact human kidney across multiple length scales without using antibodies, dyes or contrast agents. Within the human kidney, we delineated the extent and morphology of the vasculature, down to afferent and efferent arterioles, quantifying variation in vascular morphology within the context of vascular ordering schemes. This enabled quantitative comparison between human and previously published rodent kidney vascular networks, the latter of which has been used as inputs for biophysical modelling of human vascular flow [4], [5], [6], [7], [8]. We further demonstrate regional heterogeneity in the context of the anatomical compartments of the kidney. Such variations highlight the link between regional structure and function, re-enforcing the importance of quantitative analyses for understanding and modelling regional micro-environments within the human kidney.

## METHODS

### Sample preparation

An intact human kidney was obtained from a 63-year-old male (cause of death: pancreatic cancer) who consented to body donation to the Laboratoire d’Anatomie des Alpes Françaises before death. Post-mortem study was conducted according to Quality Appraisal for Cadaveric Studies scale recommendations [27]. The body was embalmed by injecting 4500 mL of 1.15% formalin in lanolin followed by 1.44% formalin into the right carotid artery, before storage at 3.6°C. During evisceration of the right kidney, vessels were exposed, and surrounding fat and connective tissue removed. The kidney was post-fixed in 4% neutral-buffered formaldehyde at room temperature for one week. The kidney was then dehydrated through an ethanol gradient over 9 days to a final equilibrium of 70% [27]. Each solution was four-fold greater than the volume of the organ and during dehydration, the solution was degassed using a diaphragm vacuum pump (Vacuubrand, MV2, 1.9m^3^/h) to remove excess dissolved gas. The dehydrated kidney was transferred to a polyethylene terephthalate jar where it was physically stabilised using a crushed agar-agar ethanol mixture, and then imaged [26], [27].

### Scanning, image acquisition and reconstruction

Imaging was performed on the BM05 beamline at the ESRF following the HiP-CT protocol [26], [27]. Initially the whole kidney was imaged at 25 µm per voxel (isotropic edge length). Volumes of interest within the same kidney were also imaged at 6.5 and 2.6 µm per voxel. [26] Tomographic reconstruction was performed [26], [27], using the PyHST2 software [28]. Briefly, a filtered back-projection algorithm with single-distance phase retrieval coupled to an unsharp mask filter was applied to the collected radiographs. Reconstruction parameters are provided in image metadata. The reconstructed volumes were binned (averaged) to 50, 13, and 5.2 µm per voxel respectively to reduce computational load for subsequent image segmentation and quantification (**see Figure S1**). All reconstructed image volumes and metadata can be accessed at human-organ-atlas.esrf.eu, table for direct DOI links for each dataset is provide in **Table S1**

### Image filtering, enhancement, and segmentation

Prior to semi-automated segmentation, images were filtered to enhance blood vessel contrast using Amira v2021.1 software. A 3D median filter (iterations =2 and 26 neighbourhood analysis) was used to reduce image noise and image normalisation was performed using background detection correction (Amira v2021.1; default parameter settings). Semi-automated segmentation of the arterial networks was performed in Amira v2021.1 using a manual region growing tool where the user selects an initial voxel within the vasculature along with set intensity and contrast thresholds. Any voxel within the connected neighbourhood of the initially selected voxel that has an intensity and contrast within the thresholds are added to the region. The annotator continues this process in an iterative fashion selecting seed points altering the thresholds to expanding the region, (Method shown in **Supplementary Video 3)**. Once the primary annotator believes they have filled the interior of all vessels the data is passed to a second annotator who repeats the process. A third annotator (referred to as the proof-reader) will then quantitatively review the labels. The data the proof-reader is presented with are randomised regions of 2D slices of the data. They then count the number of vessels cross-sections present in the slice, recoding the true positive and false negative number of vessel cross-sections that have been segmented. The proof-reader returns the data to the initial two annotators and the whole process repeats iteratively until the proof-reader does not find any false negatives. This method was applied to segment the kidney arterial network from the intact human kidney from the imaging data at 50 µm per voxel, and portions of the same network in the 13 and 5.2 µm per voxel datasets.

A second approach to quantitative validation of the segmentation was performed using smaller segmented regions of the 13 µm per voxel dataset. Here, the higher resolution volume of interest at 13 µm per voxel was rigidly registered to the whole organ volume with affine registration toolkit (Amira-Avizo) (See **Supplementary Methods §1.1 and Tables S2 & S3**). Overlapping portions of the 13 µm voxel segmentations and 50 µm per voxel datasets were extracted and the 50 µm per voxel datasets was up-sampled to the resolution of the 13 µm voxel dataset. An overlap measure known as topological precision and recall score following Paetzold et al. [29], was applied (see **Supplementary Information §2.1** & **Figure S3)**

### Visualization and skeletonization

To quantify branching metrics of the human kidney vasculature, the segmented 3D vascular network at 50 µm per voxel was skeletonized using the centreline tree algorithm in Amira-Avizo. The choice of skeletonization algorithm and the parameterising of the algorithm were optimised by utilising the super-metric approach outline by Walsh & Berg et al. [7] (tube parameters: slope = 4 and zeroval = 10, see **Supplementary Methods § 2.2 and Figure S4** for parameter selection method). The resulting spatial graph describes the vessel network in terms of ‘nodes’, ‘points’, ‘segments’, and ‘sub-segments’. A segment is defined as being between a start and end node; which correspond to either a branching point leading into another segment branch or a terminal end where no further branches were detectable. Between the start and terminal node of each segment lie sub-segments with ‘points’, marking the start and end of each sub-segment. Each sub-segment has an associated radius and length (**Figure S5A**). A multiscale smoothing approach was applied to the larger vessels (those of Strahler generation greater than 5), through a weighted smoothing algorithm, and corrections for the radius of collapsed vessels (see **Supplementary Methods § 2.3** for details). The final spatial graph was manually proofread to mitigate errors in node locations and remove spurious branches in large collapsed segments of vessels.

### Morphological analysis

Topological/morphological metrics of the network were calculated from the spatial graph as follows codes are provided at https://github.com/HiPCTProject/Skeleton_analysis:

i. branching angle, calculated as either (a) the angle between the two child segments from a common parent segment or (b) the angle between a child segment and its parent segment. In both cases the vector for the segment of parent and child were calculated between the start node and end node (i.e. ignoring vascular tortuosity);
ii. tortuosity defined as the Euclidean distance between start and end node of a segment divided by the sum of all subsegment lengths;
iii. radius calculated per segment as either, the mean of all subsegment radii, or for larger vessels that had fully collapsed (See **Supplementary Methods § 2.3** for details), as the equivalent radius for the perimeter of vessel cross-section in the binary image;
iv. length defined as the sum of all subsegment lengths;
v. inter-vessel distance calculated by two approaches to facilitate different analyses. Firstly, using the segmentation binary image the distance of every non-vessel voxel from its nearest vessel voxel was calculated *via* a 3D distance transform (ImageJ) applied to the binary vessel segmentation. Secondly, using the skeleton form, the Euclidean distance between the midpoint of every segment to its nearest-neighbouring segment midpoint was calculated.

Additionally, we also assessed vessel generation or order using two methods. Firstly, using the centripetal system known as Strahler ordering system [30], [31], [32], where the most distal segments are assigned as the first order, if two segments with the same order intersect, the resulting segment has order one greater. Alternatively, if two segments with different orders intersect, the higher order of the two is given to the resulting segment, **(Figure S5B).** Secondly, we took a centrifugal or ‘topological’ approach, starting with most proximal artery as generation one, at each branching node the generation is increased, this approach has been utilised by e.g. Pries and Secomb [33] **(Figure S5C)**.

From the ordering analyses we assessed the branching ratio (*γ*) defined as the antilog of the reciprocal for the linear fit to the plot of Stahler order (*O*) against the logarithm of the number of segments (*N*) in each order:

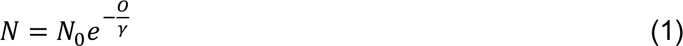

We also examined whether our data followed Murray’s Law which states that the cube of the parent segment radius should be equal to the sum of the cubed child segment radii:

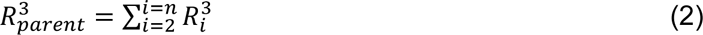

The radius of the arterial network in the human kidney obtained from this study were compared to those of the rat kidney taken from[11] scanned with 20 and 4 µm voxels using a micro-filling approach.

### Kidney Compartment Segmentation

Segmentation of the compartments within the human kidney, including cortex, medulla, inter-medullary pillars and hilum, was performed in Dragonfly (version: 2021.3) using a 2D convolutional neural network (CNN). The final hyperparameters of the CNN are given in **Table S4**. Correction of the CNN output was manually performed in Amira (v2021.1) to provide the final compartment delineation. These compartments were used to group and then analyse vascular network parameters.

### Statistical analysis

Statistical comparisons of vascular network morphology between human and rat kidneys[11] was performed in GraphPad Prism (version: 10.1.2). For all statistical tests, a *p* value of less than 0.05 was considered statistically significant. Radius against Strahler Order were normalised to the 9^th^ Strahler Order (the largest vessel for which the human data contained complete vessel segments). Log of radius against Strahler generation for each of the human and rat datasets was plotted facilitating a linear least squares regression analysis. A sum of squares *F* test was performed with the null hypothesis that a single set of global parameters for slope and intercept would fit vessel radius or vessel length for both the rat and human cases. For Murray’s law the same sum of squares *F* test was performed with the null hypothesis that Murray’s law would fit the human data; in this analysis automated outlier detection was performed with the Graphpad ROUT method where Q is the desired maximum false discovery rate, Q = 0.05%.

## RESULTS

### HiP-CT can visualise the arterial vascular network in the intact human kidney down to efferent and afferent arterioles

Using HiP-CT [26], [27] we imaged the whole intact kidney obtained from a 63-year-old male organ donor in a hierarchical fashion. We initially performed an overview scan of the entire kidney at 25 μm per voxel, followed by selecting and imaging representative volumes of interest at 6.5 μm per voxel and 2.6 μm per voxel (**Figure 1A**). As these image volumes are inherently aligned, expert annotation was applied to the image volumes taken at each resolution to produced a multi-scale segmentation of the arterial network (**Figure 1B and Supplementary Video 1**). From the segmented data we were able to identify all known anatomical subdivisions of the kidney arterial system (**Figure 1C**), down to arterioles that terminate in the specialised plasma ultrafiltration units of the kidney: the glomerulus. The segmental pattern of anterior, posterior, superior and inferior territories supplying the kidney parenchyma were clearly delineated. Each vascular territory (**Figure 1D and Supplementary Video 2**) had a corresponding kidney arterial branch originating from the hilum, which bifurcated before hierarchical branching towards the cortical parenchyma.

**Figure 1.**
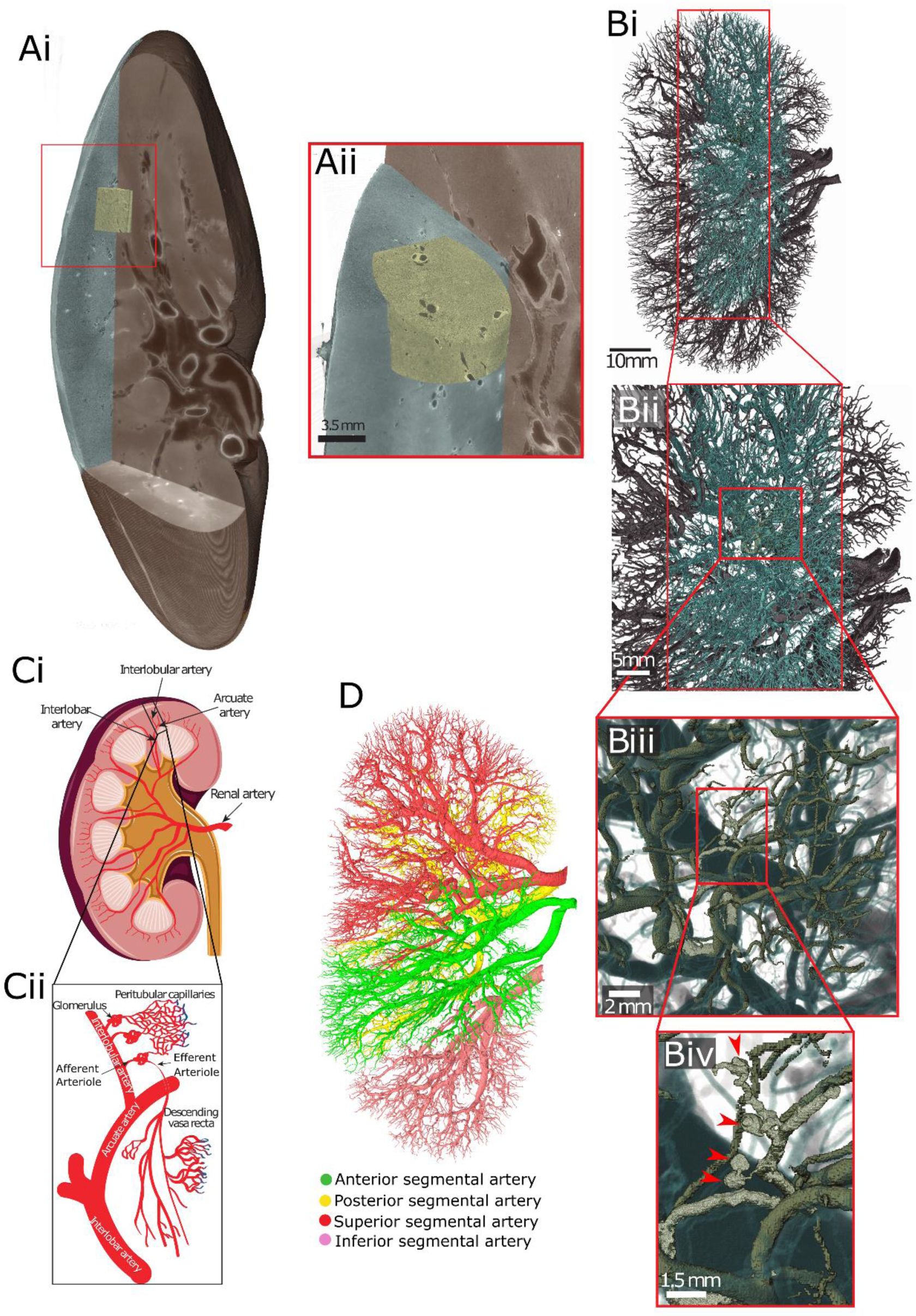
Multi-level segmentation of the human kidney arterial network. **A)** Overview of the hierarchical image volumes that can be acquired with Hierarchical Phase-Contrast Tomography (HiP-CT). Brown, cyan and yellow volumes show the whole organ acquired at 25 µm, sub-volume acquired at 6 µm and sub-volume acquired at 2.6 µm, respectively, in the intact human kidney. **B i-iv)** showing the vascular segmentation performed across the three resolutions of HiP-CT data enabling the whole organ (**Bi**) through to glomeruli (red arrows **Biv**) to be visualised and segmented. **C)** Diagram of the anatomical organisation of the human kidney arterial network. **D)** The vascular territories of the kidney imaged in this study.

### An error-bounded image processing pipeline to reproducibly quantify the arterial network in human organs

We next sought to quantitate the arterial network in a reliable and reproducible manner. As we have previously shown that quantitative features of vascular networks are influenced heavily by the image processing pipeline [7], we developed an image processing pipeline (**Figure 2**), involving reduction of the initial image to a skeleton or spatial graph representation of the arterial network. The graph representation comprises a set of nodes; 3D locations where vessels meet or end, and the connections between these nodes, defined as ‘segments’ (see **Figure S5A and Figure 4A**). Our pipeline, which is overviewed in **Figure 2**, comprises 8 steps which are fully detailed in **Supplementary Methods §2**, and enables the generation of a spatial graph from segmented HiP-CT with quantification of error at the segmentation and skeletonization stages.

**Figure 2.**
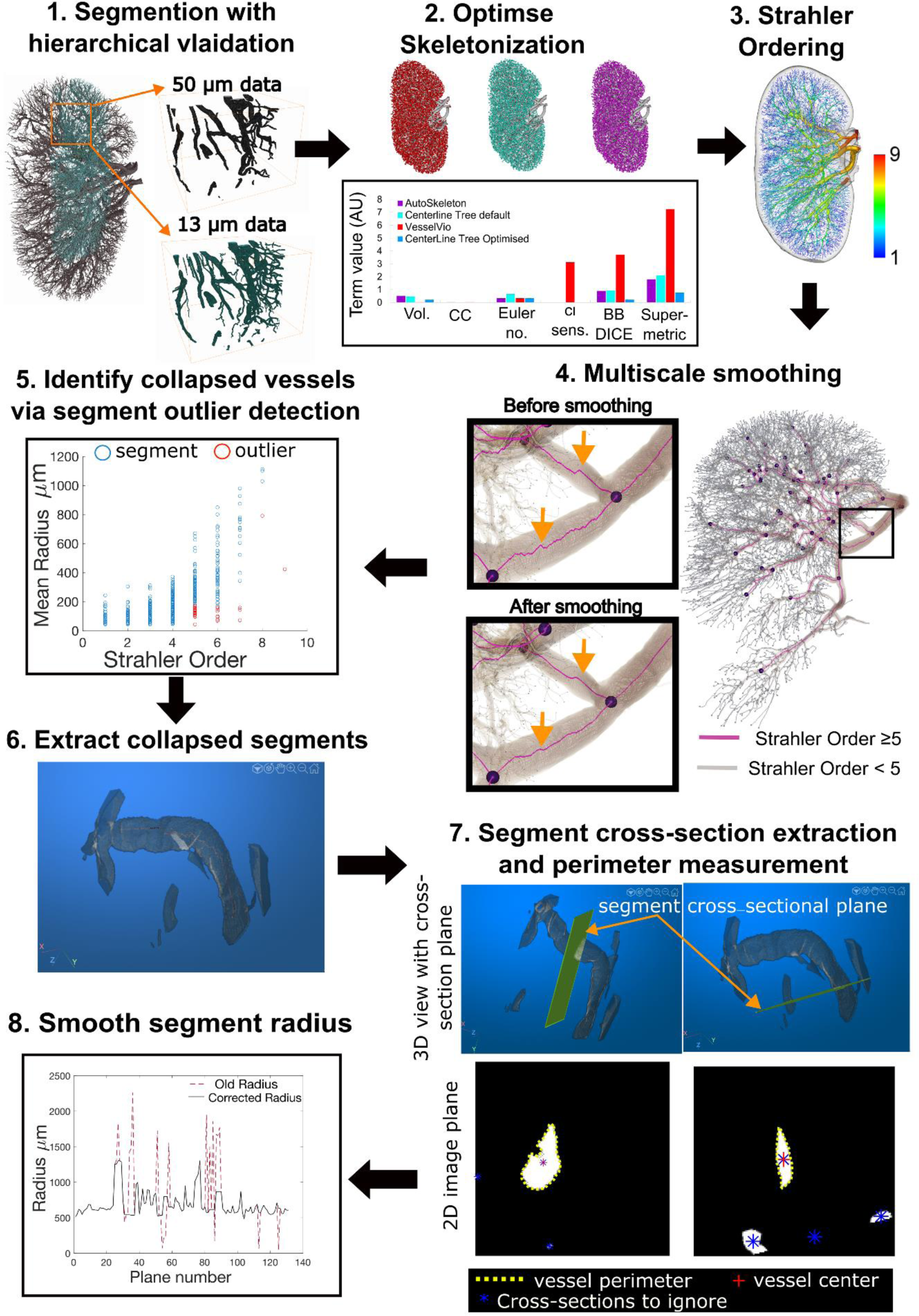
Pipeline for the extraction and correction of the vascular network skeletonization. **Step 1** Segmentation is performed with quantitative validation using a higher resolution volume of interest, **Step 2** Skeletonization is optimised by comparison of skeletonization algorithms and optimisation of skeleton super-metric, the super-metric is a projection of the distance vector between the reconstructed skeleton sand the segmented image, onto a weighted space, it contains 5 contributing terms: network volume (Vol.), connected components (CC), Euler Number, Centerline sensitivity (cl sens.), Bifurcation DICE (BB DICE). **Step 3** An initial Strahler Order (O) calculation is made on the skeletonized network. **Step 4** Using the ordering from Step 3 the network can be split into larger calibre (O ≥ 5) and smaller calibre vessels (O > 5), the larger calibre vessel can then be smoothed as shown in insets, orange arrows show the points where smoothing has noticeably acted on regions of larger vessels. **Step 5** Strahler order vs Mean radius is plotted for every segment (blue circles); outliers (red circles) are identified as segments with a radius below the 90% percentile for their Order. **Step 6** The segments identified as outliers are visualised and collapse status is manually confirmed. **Step 7** For vessels which are confirmed as collapsed, planes which are normal to the centreline of the vessel (indicated by orange arrows) are created at every point along the centreline and the 2D image for each plane is extracted (lower panels). From these 2D planes the collapsed vessel is identified (red cross) and the perimeter (yellow dashed line) is extracted. **Step 8** The perimeter is used to calculate an equivalent radius and assigned as the new radius of the segment.

In brief, the pipeline included utilising the aligned higher resolution HiP-CT volumes to provide quantification of the segmentation accuracy (**Figure 2, Step 1**). Secondly, we applied three different skeletonization algorithms and utilised a formal metric (the recently developed skeleton super-metric [7]) to assess and optimise the skeletonisation process **(Figure 2 Step 2)**. The skeleton was then corrected (**Figure 2 Steps 3 – 8**) to overcome the two primary challenges of HiP-CT data. The multiscale nature of the vasculature captured was corrected with a multi-scale smoothing approach; **Figure 2 Steps 3 and 4.** Collapsed vessels were identified from outliers in radial distributions (**Figure 2 Steps 5),** followed by automated extraction of vessel cross-section and radius correction using the collapsed vessel perimeter **Figure 2 Steps 6-8.**

The result of our novel pipeline was the generation the first open-source spatial graph of the human kidney arterial vasculature in its entirety. We were able to identify 97% of vessels <50 µm radius across the whole intact human kidney, with an imaging resolution of 50 µm per voxel. The network consisted of 10,193 nodes, 376,603 points and 10190 segments, or vessels. The total network volume was 1.68 x 10^12^ µm^3^ its was length of 2.3x10^7^ µm. This spatial graph, which is provided as spatial graphs in **Supplementary data** captures all the morphological features and connectivity of the human kidney arterial vasculature, which was then used for downstream analyses as described below.

### Multi-scale generational and ordering analysis of the arteriolar vasculature in the human kidney

Having produced a reproducible spatial graph of the human arterial vasculature of the kidney, we then performed topological generation [33] and Strahler Ordering [30], [31], [32] analyses. This resulted in nine Strahler Orders (**Figure 3A**) and twenty-five topological generations (**Figure 3B**). As the main artery supplying the kidney was cut during autopsy we can infer that 10 Strahler order, 27 topological generations, can be imaged over the entire intact human kidney with HiP-CT.

**Figure 3.**
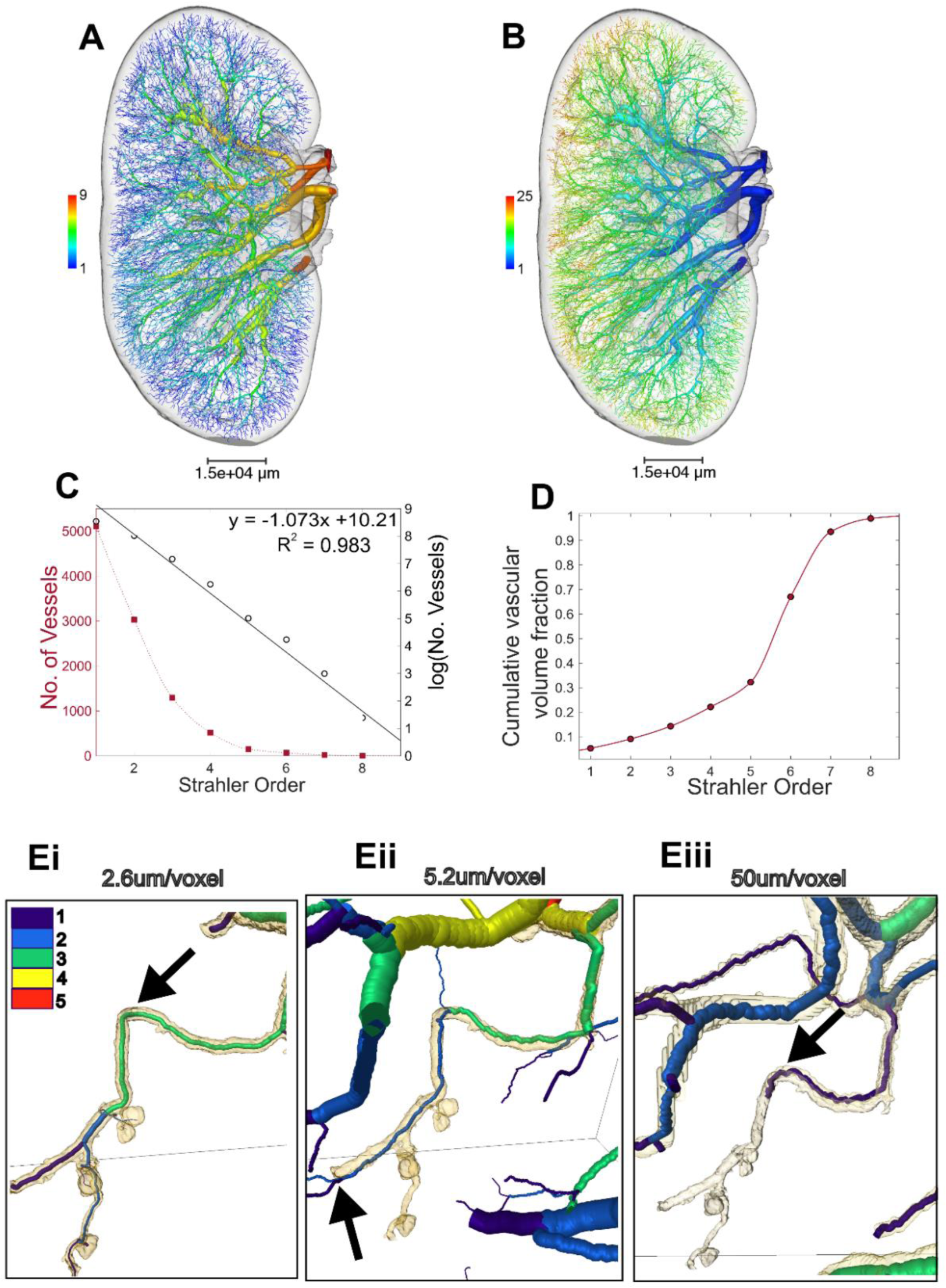
Ordering and branching ratio analyses. Rendering of the vascular network with vessels coloured according to **A)** Strahler order and **B)** Topological generation. **C)** Plot showing the number of vessels per Strahler order with fit for the log plot to calculate branching ratio. **D**) Strahler order against cumulative vascular volume fraction. **Ei**) Strahler ordering down to efferent and afferent arterioles, using the 2.6 µm/voxel image dataset; **Eii)** showing the same region segmented at 5.2 µm/voxel (binned data); and **Eiii)** the same small region from the whole kidney overview (50 µm/voxel). Black arrows in **Ei** and **Eiii** indicate the same vessel in both cases with Strahler order of 1 or 3 respectively indicated by the colour cases. The black arrow in **Eii** indicates a bifurcation downstream of the arterioles that can be segmented, which is not an arteriole itself, i.e. this demonstrates the presence of non-terminal arterioles in the human kidney.

Leveraging the hierarchical capability of HiP-CT we imaged regions of the cortex at 2.6 µm per voxel and from these data, we segmented down to the afferent arteriolar level, evidence by the presence of glomeruli at the terminal ends of these vessels **(Figure 3Ei-3Eiii)**. Interestingly, we were also able to segment glomeruli from non-terminal arteries, (the artery from which the arterioles branch have additional bifurcations downstream, which are not arterioles) (**Figure 3Eii black arrow and Figure S7**). This supports recent findings, [34] in the rat kidney, which demonstrated the existence of non-terminal branch arterioles and their contribution to the synchronicity of blood flow in the kidney [5], [35].

Glomeruli branching from non-terminal arterioles (see arrow in **Figure 3Ei** compared to **3Eiii**) prevents the application approaches used by Nordlestten et al. [11] to estimate the Strahler Order of every terminal node in the whole arterial tree (extracted from the 50 µm/voxel data) relative to glomeruli. However, our high resolution data shows there are at least 12 Strahler orders (an additional two) between afferent arterioles and the kidney artery. The estimate of 12 Strahler Orders between afferent arterioles and the kidney artery can also be supported by considering the estimates of total glomeruli number and number of terminal nodes in the 50 µm/voxel data (**See Supplementary Methods §3** and **Figure S7**). Given the scale of the arterial tree captured at our 50 µm per voxel dataset, we used this to perform further quantitative analysis.

By plotting the number of segments within each Strahler order (**Figure 3C**) we determine the branching ratio arterial vessels in the human kidney to be 2.921. This value is similar to that of the human pulmonary arterial tree (3.0 [32]) and to that of the rat kidney (2.85 [11]). To give spatial context to our data, we mapped Strahler orders to known anatomical subdivisions of the human arterial tree, including interlobar, arcuate and interlobular arteries. Strahler orders 7-9 (*n =* 25 segments; mean radius = 929 ± 477 µm) mapped to the branches of the kidney artery entering the kidney hilum. Orders 5-6 comprised interlobar arteries (*n* = 219 segments; mean radius = 417 ± 247 µm), and orders 2-4 arcuate arteries (*n* = 4841 segments; mean radius = 78 ± 45 µm). Interlobular arteries fell within orders 1-3 (*n =* 9430 segments; mean radius = 55 ± 23 µm). We further plotted the cumulative volume of the kidney vascular network is plotted in **Figure 3D**, finding that over 1/5 of the volume of the network lies within Strahler orders 1-4, corresponding to segments from interlobular arteries to arcuate arteries.

### Analysis of vascular network metrics in the human kidney reveals limit of Murray’s law and concordance with a rodent model organism

Vascular network geometric properties, such as, diameter, length and branching angles, are an important means of quantitatively comparison of vascular networks in health or disease [36], [37]. To address this, we have extracted and reported the metrics for the human kidney vasculature. We grouped our data according to Strahler order (**Figure 4**, **Table 1**) to enable quantitative comparison to rat and other human organ data, and also provide raw data for each segment in **Supplementary data** as inputs for modelling applications.

**Figure 4.**
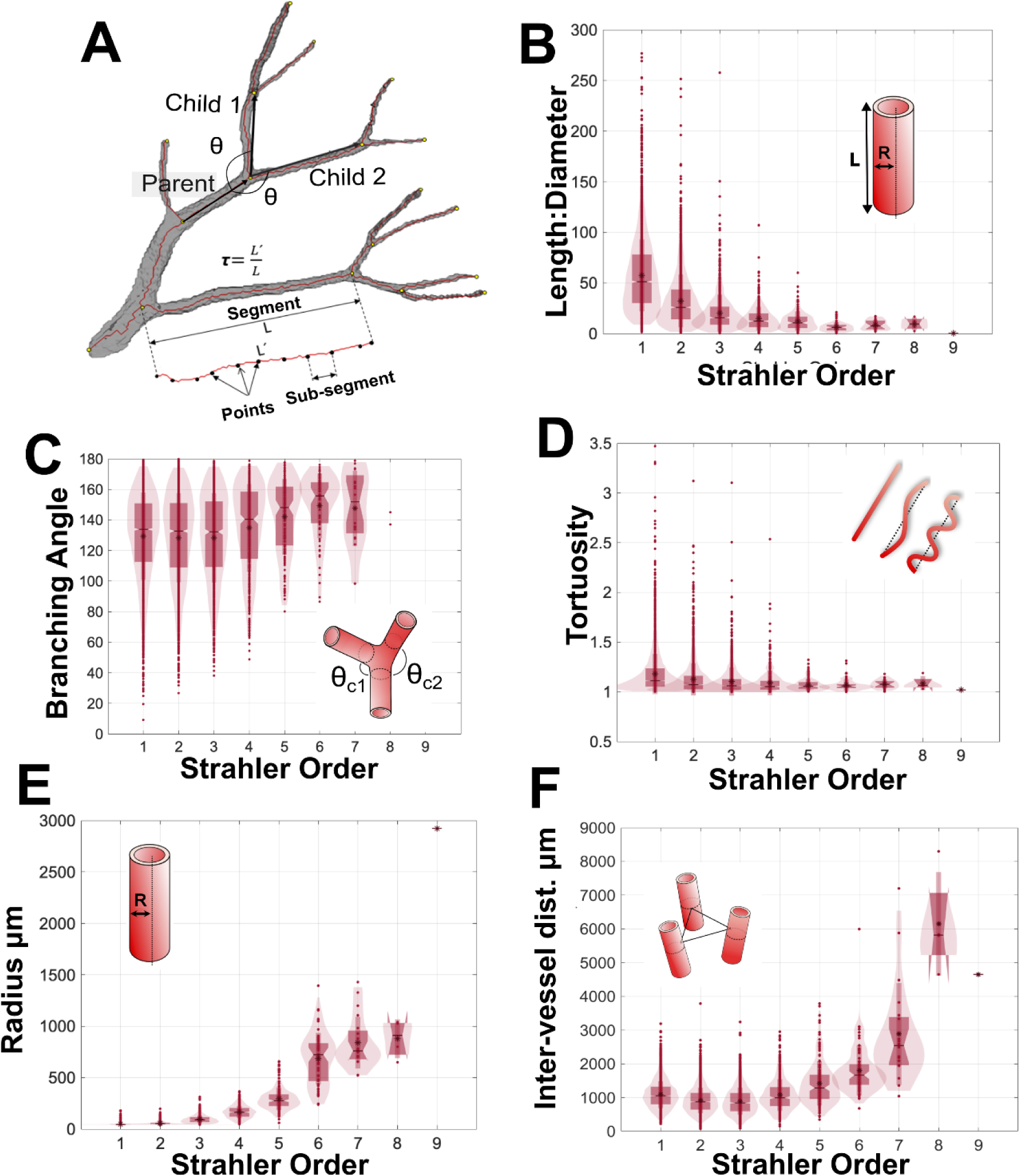
Metrics of the adult human kidney arterial network. **A)** Schematic diagram of how the metrics in B-E are calculated. **B)** The length:diameter ratio. **C)** The branching angle between the child and parent segments. **D)** The tortuosity of segments, **E)** their radius, and **F)** the inter-vessel distance as measured between the mid-point of each segment.

**Table 1.** Human kidney vascular branching metrics by Strahler generation (means with standard deviation are shown).

| Strahler order | Segments | Rad $\mu\text{m}$ | Length $\mu\text{m} \times 10^3$ | Tort. | LDR | Vol. $\times 10^8 \mu\text{m}^3$ | Branching Angle $^\circ$ | IVD $\mu\text{m} \times 10^3$ |
| --- | --- | --- | --- | --- | --- | --- | --- | --- |
| 1 | 5105 | 45±5 | 2.6±1.7 | 1.2±0.20 | 57.4±36 | 0.18±0.22 | 129±28 | 1.1±0.4 |
| 2 | 3030 | 56±15 | 1.8±1.4 | 1.1±0.16 | 32.3±26 | 0.21±0.46 | 128±29 | 0.9±0.4 |
| 3 | 1295 | 95±37 | 1.8±1.4 | 1.1±0.14 | 20.2±18 | 6.7±10.3 | 128±29 | 1.1±0.4 |
| 4 | 516 | 165±60 | 2.3±1.9 | 1.1±0.13 | 14.6±12 | 25.5±34.9 | 135±29 | 1.4±0.5 |
| 5 | 150 | 294±110 | 3.3±2.6 | 1.1±0.05 | 12.2±9.4 | 11.3±17.1 | 142±25 | 1.8±0.7 |
| 6 | 69 | 684±250 | 4.3±3.1 | 1.1±0.06 | 6.5±4.5 | 84.6±104 | 149±21 | 2.9±0.7 |
| 7 | 20 | 839±251 | 7.2±4.9 | 1.1±0.05 | 8.5±4.8 | 223±302 | 148±24 | 6.1±1.5 |
| 8 | 4 | 877±188 | 8.1±4.7 | 1.1±0.07 | 9.6±6.3 | 227±159 | 141±6 | 4.7±1.5 |
| 9 | 1 | 2923 | 669 | 1.0 | 0.2 | 180 | - | - |

As Strahler order increases, in the human kidney vascular network there is a reduction in the ratio of vessel length:diameter (**Figure 4B**), whilst the mean radius (**Figure 4E**) and inter-vessel distance increase (**Figure 4F**). Tortuosity does not vary significantly with Strahler order (**Figure 4D**); with most segments having tortuosity close to 1 implying limited deviation from a straight path. The above findings are largely consistent with anticipated trends for healthy tissue i.e. where a vasculature network is assumed to be a fractal structure, with branching pattern driven by optimised delivery of blood to the whole organ. Interestingly for Strahler orders 8-6 the mean branching angle is approximately 150°, which decreases slightly to 130° for Strahler orders 3-1 (**Figure 4C**), the latter being the predicted optimal theoretical branching angle for vascular growth that is volume constrained [38].

As simulation of kidney haemodynamics has previously been performed using micro-CT data from the rat kidney, we used our high-resolution segmentations to align our network with those derived the previously published rat dataset [11]. In doing so, we were able to relate normalised vessel metrics from each species at corresponding Strahler orders. The increase in vessel radius with Strahler order followed a similar trend between human and rat kidney (**Figure 5A**); The only parameter that showed a considerable difference was radius, as human and rat vary significantly different based on a fit of log(radius) to Strahler order (**Figure 5B)**, (p<0.0001 Sum-of-F test F (DFn, DFd) = 700.6 (2, 12)), with human vessel radii increasing with Strahler Order more quickly than the rat. To provide physiological relevance to this discrepancy, we compared our radius data to the Murray’s Law; a theoretical relationship between the radii of parent and child vessels derived from considering an optimisation of energy between blood flow through a network and diffusion into tissue with fixed metabolic demands [35], [38], [39]. The application of Murray’s Law has been previously supported by data from e.g. Nordlestten et al. which shows a deviation from Murry’s law by ∼1% for the rat kidney [11].

**Figure 5.**
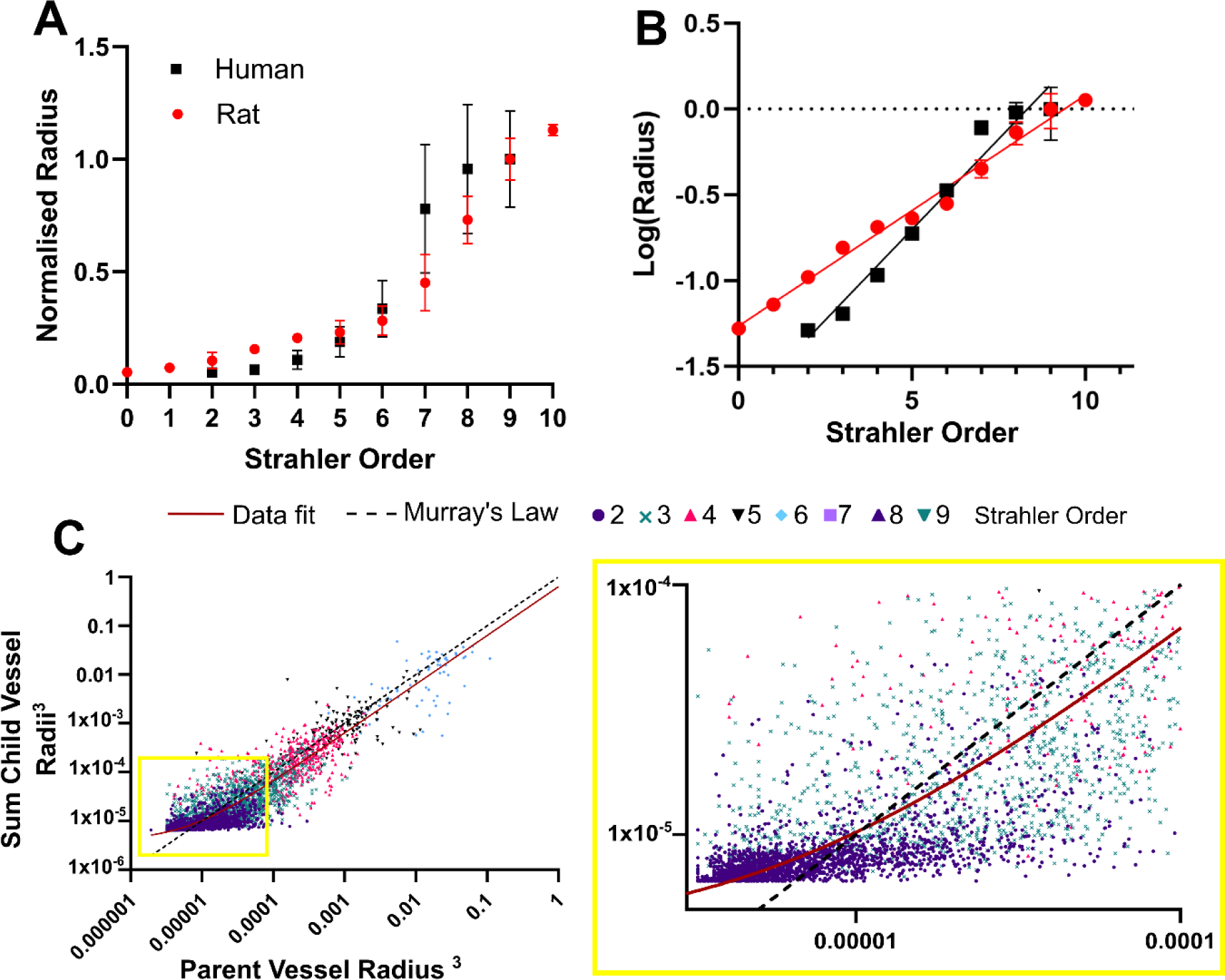
Comparison to rodent data and Murray’s theoretical law of energy balance. **A)** Normalised radius against Strahler Order for our data and for the rat data of Nordsletten et al. [11], **B)** for the same data but plotted for log(Radius), a similar pattern is seen but significant statistical difference is found between the best fit for the two datasets. **C)** Our data plotted to compare to Murray’s Law, plotted showing each Strahler order in a different colour. The best fit line for the data is shown in red with the theoretical Murray’s law in dashed black. The inset shows how the human data differs from Murray’s law predominantly for the smaller vessels.

When a best fit linear regression is applied to our data (red line **Figure 5C**), the slope 0.63 and intercept 4x10^-6^ values are with an R^2^ = 0.68, applying a Extra-sum-of-F test, F (DFn, DFd) = 5474 (1, 3949) between our data and a theoretical Murray’s law we found a significant difference P<0.0001 between Murray’s law and our data fit (**Figure 5C**). The deviation from Murray’s law is more pronounced for smaller vessels (**Figure 5C** inset). and is consistent with the previously calculated branching ratios, thus inferring deviation from a symmetric network as arterial vessels undergo hierarchal branching towards the cortex of the organ.

### Regional heterogeneity within the kidney creates local microenvironments that enable specialised kidney functions

Having demonstrated deviation in Murray’s law at lower Strahler orders, corresponding, we then sought to resolve regional heterogeneity in the human kidney vasculature. This regional heterogeneity is postulated to reflect different functions corresponding to the anatomical zone of the organ. For example, the kidney medulla possesses low oxygen tension, generating hypoxia that is inherent to the medulla’s urinary concentration mechanisms. A longstanding hypothesis, supported by blood oxygenation level-dependent MRI studies, [40] is that vascular rarefaction in CKD results in hypoxia within the kidney cortex, stimulating neighbouring cells into a pro-fibrotic phenotype and manifesting in loss of organ function [3]. Although the regional heterogeneity of vascular patterning is likely to be fundamental for local microenvironments, such as the generation of physiological hypoxia or susceptibility to pathological hypoxia, it has not been quantitatively explored in the human kidney vasculature.

Leveraging the contrast-free approach of HiP-CT imaging, we were able to segment the kidney into known anatomical compartments including hilum, medulla, intramedullary kidney columns and cortex (**Figure 6A**). The total tissue volume of each zone in addition to the number of vessels, length, radius and volume of segmented vessels within each zone were quantified (**Table. 2**). Most of the tissue volume of the human kidney was measured to be occupied by the cortex (63.7%) as compared with the medulla (23.5%) or hilum (8.7%) or intermedullary pillars (4.1%). The number of segments for the vascular network within each compartment followed this trend. As a proxy for kidney tissue oxygenation, we quantified (**Figure 6Ai-Aiii**) and mapped (**Figure 6Bi-Biii**) the inter-vessel distance, reflecting the extravascular distance across which oxygen and solutes diffuse, compartmentalised by hilum, medulla, cortex, and inter-medullary pillars. Mean inter-vessel distances for these compartments **(Table. 2)** demonstrates the medulla having the highest inter-vessel distance which follows the anticipated distributions of hypoxia. Notably, there were large portions within the medulla where inter-vessel distance was > 4.5 mm **(Figure 6Bi)**; in line with the known hypoxic characteristic of the medulla. Whilst the cortex has one of smallest inter-vessel distance, it has a large standard deviation and the heatmap in **Figure 6Bi and 6Bii** shows small areas with inter-vessel distance > 4.5mm are also found, predominantly towards the kidney capsule.

**Figure 6.**
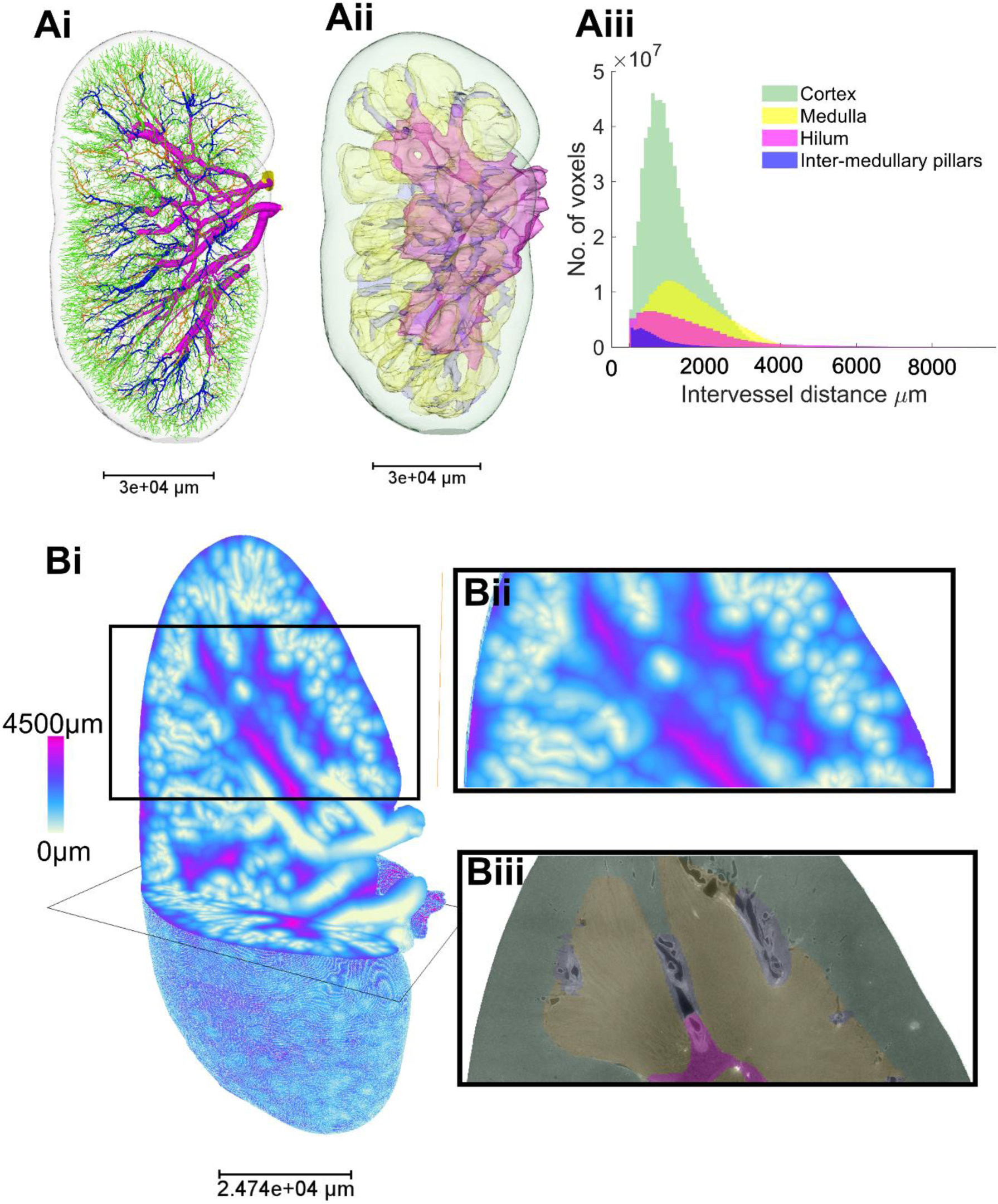
Analysis of zonal heterogeneity in vascular branching metrics within the human kidney. **(Ai)** 3D reconstruction vasculature colour according to anatomical compartment within the human kidney cortex (green), medulla (yellow) and hilum (pink), inter-medullar pillars (dark blue). (**Aii**) Showing the 3D surface masks of the same regions. **(Aiii)** Inter-vessel distances are plotted against the total number of vessel voxels for each kidney compartment. **(Bi)** Visual heatmap of inter-vessel distance for the entire human kidney, where pink represents the largest inter-vessel distance (> 4.5 mm) and white (0 mm) the smallest. (**Bii**) A digital zoomed region within cortex and medulla. **(Biii)** The 2D slice of the associated HiP-CT raw image with the compartments overlaid.

**Table 2.** Human kidney vascular branching metrics by zone.

|  | Cortex | Medulla | Hilum | Inter-medullary pillars | Organ |
| --- | --- | --- | --- | --- | --- |
| Volume of tissue $\times 10^{13} \mu\text{m}^3$ (% of total) | 8.70 (63.7%) | 3.21 (23.5%) | 1.18 (8.66%) | 0.57 (4.14%) | 13.7 (100%) |
| Number of segments* (% of total) | 6141 (60.27%) | 554 (5.4%) | 151 (1.5%) | 727 (7.1%) | 10190 (100%) |
| Mean segment length, $\mu\text{m} \pm \text{STD}$ | 1999±1374 | 1493±1113 | 3993±3568 | 1720±1386 | 2260±1720 |
| Mean segment radius, $\mu\text{m} \pm \text{STD}$ | 48±12.6 | 95±49 | 496±335 | 136±80 | 71±87 |
| Mean inter-vessel distances, $\times 10^3 \mu\text{m} \pm \text{STD}$ | 1.10±0.677 | 1.55±0.881 | 1.55±1.312 | 0.664±0.543 | 1.2±0.833 |
| Mean segment volume, $\times 10^8 \mu\text{m}^3 \pm \text{STD}$ | 0.148±0.116 | 0.623±1.12 | 69.0±147 | 1.73±4.48 | 1.65± 20.6 |
| Mean segment tortuosity $\pm \text{STD}$ | 1.14±0.17 | 1.08±0.13 | 1.08±0.1 | 1.1±0.12 | 1.15±0.18 |
| *segments that crossed over two regions were excluded. |  |  |  |  |  |

## DISCUSSION

Owing to the limited volume of tissue imageable using pre-clinical modalities such as micro-CT and lightsheet microscopy, and insufficient resolution of technologies routinely used in clinical practice such a CT and MRI, it had previously been impossible to capture the entire vascular network of the intact adult human kidney. Using synchrotron-based HiP-CT we were able to segment and quantify the human kidney arterial network from kidney artery to interlobular arteries without the need for exogenous contrast agents. With this method, we demonstrated that vessels mapping from interlobar to interlobular arteries occupy approximately 1/5^th^ of the vascular volume of the human kidney. By imaging regions of interest in the intact kidney at higher resolution and aligning with low resolution data, we further demonstrate that, akin to rat [34] and varying from the traditional hierarchy of the kidney vasculature observed in textbooks, that the glomeruli in humans can originate from non-terminal arterioles. Although there existed similarity in topology between human and rat kidney, there was a significant difference in the change in radius with Strahler order between organisms. As vessels branch towards the cortex and radius decreases in the human kidney vasculature, they also do not adhere to Murray’s theoretical law of energy balance. Finally, we demonstrate vascular volume fractions and inter vessel distances vary between anatomical zones of the kidney, corresponding to regionally specialised functions and known physiological gradients in local oxygen tension.

The human kidney vasculature is exquisitely specialised to meet the physiological demands of the kidney. Underpinning this specialisation is the cellular and molecular heterogeneity of endothelial beds with the renal vasculature [41] of which we are gaining an increasing understanding due to the advent of improved techniques such as cellular transcriptomics. The rapid advances in our understanding of cellular and molecular heterogeneity of the kidney vasculature has not been matched by structural insights, likely as a consequence of limitations in imaging technologies. We have overcome these limitations using HiP-CT, and capture the 3D vascular architecture of an entire human kidney at twenty-fold greater resolution than conventional hospital CT scanners (400 µm per voxel), instead comparable to that of light microscopy (1-8 µm per voxel) yet on a volume many orders of magnitude larger than that of a kidney punch biopsy. The balance between imaging volume and resolution afforded by HiP-CT thus bridges the scale between local cellular architecture and global tissue structure, providing quantitative vascular branching metrics from an intact human organ for the first time. Testament to this balance, we report up to 10 Strahler orders or 27 topological generations for the human kidney vasculature. These exceed previous *in vivo MRI* studies, which report up to six topological orders [23] Despite being taken from the lowest resolution HiP-CT scan, these metrics also exceeds studies on cadaveric cast and dye injections which report up to arterial branches corresponding to Strahler orders 7-9) [22], [42]. We demonstrate concordance in topological layout between the human kidney vasculature and that of the rat, the latter of which has been key for inputs to generate biophysical models of kidney haemodynamics [5], [11] However, we demonstrate deviations in the progressive decrease in magnitude of vessel radii between human and rat kidney vasculature, and non-adherence to a symmetrical network that would be consistent with Murray’s law. This discrepancy could potentially influence the prior simulations of haemodynamics, oxygenation or drug delivery [8], [43]; and generation of synthetic vessel trees for *in silico* experiments [44], [45]. This divergence between theoretical prediction and our human data may be accounted for by regional heterogeneity in architecture, corroborated by our segmentation of hilar, medullary, intra-medullary and cortical zones of HiP-CT images from the same kidney. The increased inter-vessel distance observed within the medullary compared to other kidney anatomical zones corresponds to the known decreased oxygen tension within this region of the kidney. Thus, structural heterogeneity could impart varying metabolic demands and generate local microenvironments within the kidney [46], [47], which may underpin the unique cellular and molecular adaptations of specialised endothelia across the kidney vascular network [41].

The quantitative analysis pipeline performed in this paper serves multiple purposes. Firstly, it surmounts one of the limitations of HiP-CT, in the size of the datasets generated. Whereas the whole kidney dataset amounts to ∼500GB, necessitating considerable computational power and data storage facilities, segmentation followed by skeletonization allows the whole kidney vasculature dataset to be represented in a spatial graph comprising only KB of data. The spatial graph, which is provided as **Supplementary Data**, is readily quantifiable. Whereas prior simulations of kidney haemodynamics and perfusion have relied on seminal micro-CT studies performed in rat, we provide, for the first time, a complete map of the kidney arterial network in its entirety. The segmentation is accurate, with 97% of vessels of <50 µm radius captured across the intact human kidney, and thus provides vital inputs for future biophysical modelling frameworks of kidney physiology. It also serves as a reference dataset to study kidney diseases, in which vascular rarefaction is a pathophysiological hallmark[48]. The pipeline could be used to generate vascular maps from multiple kidneys or other human organs, potentially giving rise to spatial ‘atlases’ of human organ vasculature across healthy and pathological contexts. Beyond these, our openly available dataset has immediate practical applications, such as providing inputs for bioprinting for tissue engineering of artificial kidneys [49] or planning surgical resection of kidney tumours whilst preserving kidney function. [50] These datasets can also be used as a tool for medical education and training, as well as for the creation and advancement of surgical methods.

There are several limitations of this work, including the low throughput of the segmentation of this type of data. Here we present the complete analysis from a single kidney, as a framework for future studies to study further kidneys in health and disease, or other intact human organs. The accuracy of the segmentation, however, lays a foundation for tools such as machine learning methods for automated segmentation of blood vasculature from imaging data,[17], [51], [52]. Although the resolution of the organ-wide HiP-CT scan far exceeds conventional *in vivo* and *ex vivo* imaging measures, it still cannot achieve arteriole resolution across the whole organ. Nevertheless, improvements of the ESRF beamline (BM18) have already been developed, and will extend the resolution limit for whole organs down to 8um, whilst increasing the speed of scanning. Although the physical access to HiP-CT via the synchrotron is limited, we have released all our data thus far through the Human Organ Atlas portal (https://human-organ-atlas.esrf.eu/), for open-access download and use of the data for biomedical researchers.

In summary, we have achieved quantitative mapping of the entire arterial network of an intact human kidney for the first time: a vital step towards understanding how physical properties of the kidney vasculature relate to cellular and molecular heterogeneity, whilst generating key inputs for future biophysical modelling of kidney vascular physiology. Ultimately, we envisage that mapping of microstructural detail will become routine at the scale of the whole kidney, providing a means to link cellular events with organ physiology and pathology.

## FUNDING

This [publication, dataset, software etc.] has been made possible in part by grants DAF2020-225394 and 2022-316777 (DOI /10.37921/331542rbsqvn) from the Chan Zuckerberg Initiative DAF, an advised fund of Silicon Valley Community Foundation, and grant CZIF2021-006424 from the Chan Zuckerberg Initiative Foundation, the MRC (MR/R025673/1), the RAEng (CiET1819-10), and ESRF beamtimes (md1252 & md1290). DALs laboratory is supported by a Wellcome Trust Investigator Award (220895/Z/20/Z) and the NIHR Biomedical Research Centre at Great Ormond Street Hospital for Children NHS Foundation Trust and University College London.

## ACKNOWLEDGEMENTS

……….

## AUTHOR CONTRIBUTIONS

P.D.L, R.J.S., and, P.T. conceptualized the project and designed experiments; P.T. designed and built instrumentation for HiP-CT imaging; P.T. J.B and C.L.W. designed and implemented tomographic reconstruction areas; S.R., J.B, S.N and C.L.W. designed, managed and performed image analysis; S.R. and C.L.W. modelled, quantified and provided results; S.R., C.L.W., R.J.S., D.A.L., D.J.J., and P.D.L, provided results interpretation and discussion; S.R., C.L.W., D.J.J., and D.A. L. wrote the paper; D.J.J. and D.A.L. provided medical interpretation for kidney analysis. All authors assisted in reviewing and revising the manuscript.

## DATA SHARING STATEMENT

The image data that form the basis of the study findings are freely available at the ESRF data repository (https://human-organ-atlas.esrf.eu). Additionally, the spatial graph data of the kidney arterial network, along with the computed morphological parameters, can be accessed in Supplementary data which will be released following peer review.

## Notes

https://human-organ-atlas.esrf.eu/

### Competing Interest Statement

ACKNOWLEDGEMENTS
The authors would like to express their gratitude for the financial support provided by the Chan Zuckerberg Initiative DAF (2020-225394), an advised fund of SVCF, the MRC (MR/R025673/1), and ESRF beamtimes (md1252 & md1290). P.D.L. is supported by a Royal Academy of Engineering Chair in Emerging Technologies (CiET1819/10). D.J.J. is supported by a Foulkes Foundation Postdoctoral Fellowship. D.A.L. is supported by a Wellcome Trust Investigator Award (220895/Z/20/Z) and by the National Institute for Health Research (NIHR) Biomedical Research Centre at Great Ormond Street Hospital for Children NHS Foundation Trust and University College London. This research was funded in part by the Wellcome Trust [209553/Z/17/Z]. M.A. is supported by the National Institutes of Health (NIH) (HL94567 and HL134229). JJ, was also supported by the NIHR UCLH Biomedical Research Centre, UK.
DISCLOSURE
JJ reports fees from Boehringer Ingelheim, Roche, NHSX, Takeda and GlaxoSmithKline unrelated to the submitted work. JJ was supported by Wellcome Trust Clinical Research Career Development Fellowship 209553/Z/17/Z and the NIHR Biomedical Research Centre at University College London.

### Summary of Updates

The data and analyses have been updated

## REFERENCES

[1] G. Molema and W. C. Aird, ‘Vascular heterogeneity in the kidney’, in Seminars in nephrology, Elsevier, 2012, pp. 145–155.

[2] S. J. Dumas et al., ‘Phenotypic diversity and metabolic specialization of renal endothelial cells’, Nat Rev Nephrol, vol. 17, no. 7, pp. 441–464, 2021.

[3] D. A. Long, J. T. Norman, and L. G. Fine, ‘Restoring the renal microvasculature to treat chronic kidney disease’, Nat Rev Nephrol, vol. 8, no. 4, pp. 244–250, 2012.

[4] L. F. M. Cury, G. D. Maso Talou, M. Younes-Ibrahim, and P. J. Blanco, ‘Parallel generation of extensive vascular networks with application to an archetypal human kidney model’, R Soc Open Sci, vol. 8, no. 12, Dec. 2021, doi: 10.1098/rsos.210973.

[5] D. J. Marsh, D. D. Postnov, O. V Sosnovtseva, and N.-H. Holstein-Rathlou, ‘The nephron-arterial network and its interactions’, American Journal of Physiology-Renal Physiology, vol. 316, no. 5, pp. F769–F784, 2019.

[6] A. d’Esposito et al., ‘Computational fluid dynamics with imaging of cleared tissue and of in vivo perfusion predicts drug uptake and treatment responses in tumours’, Nat Biomed Eng, vol. 2, no. 10, pp. 773–787, 2018, doi: 10.1038/s41551-018-0306-y.

[7] C. L. Walsh, M. Berg, H. West, N. A. Holroyd, S. Walker-Samuel, and R. J. Shipley, ‘Reconstructing microvascular network skeletons from 3D images: what is the ground truth?’, Comput Biol Med, vol. 171, p. 108140, 2024.

[8] P. Xu et al., ‘A hybrid approach to full-scale reconstruction of renal arterial network’, Sci Rep, vol. 13, no. 1, p. 7569, 2023.

[9] D. J. Marsh, D. D. Postnov, D. J. Rowland, A. S. Wexler, O. V Sosnovtseva, and N.-H. Holstein-Rathlou, ‘Architecture of the rat nephron-arterial network: analysis with micro-computed tomography’, American Journal of Physiology-Renal Physiology, vol. 313, no. 2, pp. F351–F360, 2017.

[10] D. S. Perrien et al., ‘Novel methods for microCT-based analyses of vasculature in the renal cortex reveal a loss of perfusable arterioles and glomeruli in eNOS-/-mice’, BMC Nephrol, vol. 17, no. 1, pp. 1–10, 2016.

[11] D. A. Nordsletten, S. Blackett, M. D. Bentley, E. L. Ritman, and N. P. Smith, ‘Structural morphology of renal vasculature’, Am J Physiol Heart Circ Physiol, vol. 291, no. 1, 2006, doi: 10.1152/ajpheart.00814.2005.

[12] N. Parvin, J. R. Charlton, E. J. Baldelomar, J. J. Derakhshan, and K. M. Bennett, ‘Mapping vascular and glomerular pathology in a rabbit model of neonatal acute kidney injury using MRI’, Anat Rec, vol. 303, no. 10, pp. 2716–2728, 2020.

[13] J. Foiret, H. Zhang, T. Ilovitsh, L. Mahakian, S. Tam, and K. W. Ferrara, ‘Ultrasound localization microscopy to image and assess microvasculature in a rat kidney’, Sci Rep, vol. 7, no. 1, p. 13662, 2017.

[14] J. Huang et al., ‘A cationic near infrared fluorescent agent and ethyl-cinnamate tissue clearing protocol for vascular staining and imaging’, Sci Rep, vol. 9, no. 1, p. 521, 2019.

[15] A. Klingberg et al., ‘Fully automated evaluation of total glomerular number and capillary tuft size in nephritic kidneys using lightsheet microscopy’, Journal of the American Society of Nephrology, vol. 28, no. 2, pp. 452–459, 2017.

[16] O. Ogunlade et al., ‘In vivo three-dimensional photoacoustic imaging of the renal vasculature in preclinical rodent models’, American Journal of Physiology-Renal Physiology, vol. 314, no. 6, pp. F1145–F1153, 2018.

[17] W. Zheng et al., ‘Deep Learning Enhanced Volumetric Photoacoustic Imaging of Vasculature in Human’, Advanced Science, vol. 10, no. 29, p. 2301277, Oct. 2023, doi: 10.1002/advs.202301277.

[18] S. Balıkçı Dorotea, T. Banzato, L. Bellini, B. Contiero, and A. Zotti, ‘Kidney Measures in the Domestic Rat: A Radiographic Study and a Comparison to Ultrasonographic Reference Values’, J Exot Pet Med, vol. 25, no. 2, pp. 157–162, 2016, doi: 10.1053/j.jepm.2016.03.011.

[19] M. J. Musa and A. Abukonna, ‘Sonographic measurement of renal size in normal high altitude populations’, J Radiat Res Appl Sci, vol. 10, no. 3, pp. 178–182, 2017, doi: 10.1016/j.jrras.2017.04.004.

[20] O. K. Zenin, E. S. Kafarov, O. A. Beshulya, L. A. Udochkina, and H. M. Bataev, ‘Quantitative Anatomy of the Intrainganic Arterial Kidney’, in International Conference on Health and Well-Being in Modern Society (ICHW 2019), Atlantis Press, 2019, pp. 129–132.

[21] S. Zhao et al., ‘Cellular and molecular probing of intact human organs’, Cell, vol. 180, no. 4, pp. 796–812, 2020.

[22] I. U. Vagabov, E. S. Kafarov, O. K. Zenin, T. S. Dokaeva, and K. M. Bataev, ‘Segmental Arteries as Sources of Formation of Arterial Segments of Human Kidney’, in The International Conference “Health and wellbeing in modern society”(ICHW 2020), Atlantis Press, 2020, pp. 341–346.

[23] L. Timms et al., ‘Ferumoxytol-enhanced ultrashort TE MRA and quantitative morphometry of the human kidney vasculature’, Abdominal Radiology, vol. 46, pp. 3288–3300, 2021.

[24] J. R. Charlton et al., ‘Image analysis techniques to map pyramids, pyramid structure, glomerular distribution, and pathology in the intact human kidney from 3-D MRI’, American Journal of Physiology-Renal Physiology, vol. 321, no. 3, pp. F293–F304, 2021.

[25] D. J. Jafree, et al., ‘Three-dimensional imaging and single-cell transcriptomics of the human kidney implicate perturbation of lymphatics in alloimmunity’, bioRxiv, p. 2022.10.28.514222, Jan. 2022, doi: 10.1101/2022.10.28.514222.

[26] C. L. Walsh et al., ‘Imaging intact human organs with local resolution of cellular structures using hierarchical phase-contrast tomography’, Nat Methods, vol. 18, no. 12, pp. 1532–1541, 2021.

[27] J. Brunet et al., ‘Preparation of large biological samples for high-resolution, hierarchical, multi-modal imaging’, bioRxiv, 2022.

[28] A. Mirone, E. Brun, E. Gouillart, P. Tafforeau, and J. Kieffer, ‘The PyHST2 hybrid distributed code for high speed tomographic reconstruction with iterative reconstruction and a priori knowledge capabilities’, Nucl Instrum Methods Phys Res B, vol. 324, pp. 41–48, 2014.

[29] J. C. Paetzold et al., ‘clDice—A novel connectivity-preserving loss function for vessel segmentation’, in Medical Imaging Meets NeurIPS 2019 Workshop, 2019.

[30] G. S. Kassab, K. Imoto, F. C. White, C. A. Rider, Y.-C. Fung, and C. M. Bloor, ‘Coronary arterial tree remodeling in right ventricular hypertrophy’, American Journal of Physiology-Heart and Circulatory Physiology, vol. 265, no. 1, pp. H366–H375, 1993.

[31] A. N. Strahler, ‘Quantitative analysis of watershed geomorphology’, *Eos*, Transactions American Geophysical Union, vol. 38, no. 6, pp. 913–920, Dec. 1957, doi: 10.1029/TR038i006p00913.

[32] K. Horsfield, ‘Morphometry of the small pulmonary arteries in man.’, Circ Res, vol. 42, no. 5, pp. 593–597, 1978.

[33] A. R. Pries and T. W. Secomb, ‘Blood Flow in Microvascular Networks’, Microcirculation, pp. 3–36, 2008, doi: 10.1016/B978-0-12-374530-9.00001-2.

[34] D. J. Marsh, D. D. Postnov, D. J. Rowland, A. S. Wexler, O. V Sosnovtseva, and N.-H. Holstein-Rathlou, ‘Architecture of the rat nephron-arterial network: analysis with micro-computed tomography’, Am J Physiol Renal Physiol, vol. 313, no. 2, pp. F351– F360, Aug. 2017, doi: 10.1152/ajprenal.00092.2017.

[35] D. D. Postnov et al., ‘Modeling of Kidney Hemodynamics: Probability-Based Topology of an Arterial Network’, PLoS Comput Biol, vol. 12, no. 7, pp. e1004922–e1004922, Jul. 2016, doi: 10.1371/journal.pcbi.1004922.

[36] C. O’Connor, E. Brady, Y. Zheng, E. Moore, and K. R. Stevens, ‘Engineering the multiscale complexity of vascular networks’, Nat Rev Mater, vol. 7, no. 9, pp. 702– 716, 2022.

[37] J. Ehling et al., ‘Quantitative micro-computed tomography imaging of vascular dysfunction in progressive kidney diseases’, Journal of the American Society of Nephrology, vol. 27, no. 2, pp. 520–532, 2016.

[38] E. Tekin, D. Hunt, M. G. Newberry, and V. M. Savage, ‘Do Vascular Networks Branch Optimally or Randomly across Spatial Scales?’, PLoS Comput Biol, vol. 12, no. 11, pp. e1005223–e1005223, Nov. 2016, doi: 10.1371/journal.pcbi.1005223.

[39] W. Deng and K. Tsubota, ‘Numerical simulation of the vascular structure dependence of blood flow in the kidney’, Med Eng Phys, vol. 104, p. 103809, 2022.

[40] M. Pruijm et al., ‘Renal blood oxygenation level-dependent magnetic resonance imaging to measure renal tissue oxygenation: a statement paper and systematic review’, Nephrology Dialysis Transplantation, vol. 33, no. suppl_2, pp. ii22–ii28, 2018.

[41] S. J. Dumas et al., ‘Phenotypic diversity and metabolic specialization of renal endothelial cells’, Nat Rev Nephrol, vol. 17, no. 7, pp. 441–464, 2021, doi: 10.1038/s41581-021-00411-9.

[42] M. M. Shoja et al., ‘Peri-hilar branching patterns and morphologies of the renal artery: a review and anatomical study’, Surgical and Radiologic Anatomy, vol. 30, pp. 375– 382, 2008.

[43] A. d’Esposito et al., ‘Computational fluid dynamics with imaging of cleared tissue and of in vivo perfusion predicts drug uptake and treatment responses in tumours’, Nat Biomed Eng, vol. 2, no. 10, pp. 773–787, 2018.

[44] L. F. M. Cury, G. D. Maso Talou, M. Younes-Ibrahim, and P. J. Blanco, ‘Parallel generation of extensive vascular networks with application to an archetypal human kidney model’, R Soc Open Sci, vol. 8, no. 12, p. 210973, 2021.

[45] E. Brown, et al., ‘Physics-informed deep generative learning for quantitative assessment of the retina’, bioRxiv, p. 2023.07.10.548427, Jan. 2023, doi: 10.1101/2023.07.10.548427.

[46] A. B. Brummer et al., ‘Branching principles of animal and plant networks identified by combining extensive data, machine learning and modelling’, J R Soc Interface, vol. 18, no. 174, p. 20200624, 2021.

[47] E. Tekin, D. Hunt, M. G. Newberry, and V. M. Savage, ‘Do vascular networks branch optimally or randomly across spatial scales?’, PLoS Comput Biol, vol. 12, no. 11, p. e1005223, 2016.

[48] D. A. Long, J. T. Norman, and L. G. Fine, ‘Restoring the renal microvasculature to treat chronic kidney disease’, Nat Rev Nephrol, vol. 8, no. 4, pp. 244–250, 2012, doi: 10.1038/nrneph.2011.219.

[49] C. G. Lebedenko and I. A. Banerjee, ‘Enhancing Kidney vasculature in tissue engineering—Current trends and approaches: A Review’, Biomimetics, vol. 6, no. 2, p. 40, 2021.

[50] J. J. Hsieh et al., ‘Renal cell carcinoma’, Nat Rev Dis Primers, vol. 3, no. 1, pp. 1–19, 2017.

[51] C. Kirst et al., ‘Mapping the fine-scale organization and plasticity of the brain vasculature’, Cell, vol. 180, no. 4, pp. 780–795, 2020.

[52] M. I. Todorov et al., ‘Machine learning analysis of whole mouse brain vasculature’, Nat Methods, vol. 17, no. 4, pp. 442–449, 2020.

