## Supplementary Methods for "Mapping the blood vasculature in an intact human kidney using hierarchical phase-contrast tomography"

**Running title: Analysis of the vasculature in an intact human kidney**

Shahrokh Rahmani [1,2] , Daniyal J Jafree [3, 4, 5], Peter D Lee [1], Paul Tafforeau [6],Joseph Brunet[1,6] Sonal Nandanwar[1], Joseph Jacob [7,8], Alexandre Bellier [9], Maximilian Ackermann [10,11, 12] Danny D Jonigk [12,13], Rebecca J Shipley [1], David A Long [3,5], Claire L Walsh [1]

1. Department of Mechanical Engineering, University College London, London, UK, WC1E 6BT.
2. National Heart & Lung Institute, Faculty of Medicine, Imperial College London, London, United Kingdom
3. Developmental Biology and Cancer Research & Teaching Department, UCL Great Ormond Street Institute of Child Health, University College London, London, UK, WC1N 1EH.
4. UCL MB/PhD Programme, Faculty of Medical Science, University College London, London, UK, WC1E 6BT.
5. UCL Centre of Kidney and Bladder Health, UCL London UK
6. European Synchrotron Radiation Facility, Grenoble, France, 38043.
7. Satsuma Lab, Centre for Medical Image Computing, UCL, London, UK.
8. Lungs for Living Research Centre, UCL, London, UK.
9. Department of Anatomy (LADAF), Grenoble Alpes University, Grenoble, France, 38058.
10. Institute of Anatomy, University Medical Center of the Johannes Gutenberg University Mainz, Mainz, Germany.
11. Institute of Pathology and Department of Molecular Pathology, Helios University Clinic Wuppertal, University of Witten-Herdecke, Wuppertal, Germany.
12. Institute of Pathology, RWTH Aachen Medical University, Aachen, Germany.
13. German Center for Lung Research (DZL), Biomedical Research in Endstage and Obstructive Lung Disease Hannover (BREATH), Hannover, Germany.

### 1 Acquisition and Reconstruction Image

Reconstructed datasets can be obtained and downloaded from **human-organ-atlas.esrf.eu** where they can also be viewed in-browser through neuroglancer. Metadata for each dataset including scanning parameters, demographic and medical donor information and reconstruction parameters are available through this portal and each dataset can be accessed and cited via a separate DOI as listed in **Table S1**.

| Dataset name | Voxel size (µm) | DOI |
|---|---|---|
| 25.0um_LADAF-2021-17_kidney-right_complete-organ | 25 | DOI10.15151/ESRF-DC-1773966439 |
| 6.5um_LADAF-2021-17_kidney-right_VOI-03 | 6.5 | Will be updated on Publication |
| 2.6um_LADAF-2021-17_kidney-right_VOI-03.1 | 2.6 | Will be updated on Publication |

**Table S1 Dataset availability.**

#### 1.1 Registration of local zoom scans

The higher resolution volumes of interest (6.5 µm (down sampled to 13 µm) and 2.6 µm (downsampled to 5.2 µm)) were registered to the 25 µm/voxel (downsampled to 50 µm/voxel) image volume to facilitate annotation of vasculature across the scales. Image volumes were registered in in Amira-Avizo V2022.1 using mutual information metric, with rigid alignment and isotropic constraints. The initialisation of the volume positions was provided manually by the annotator. Parameter options are shown in **Figure S2**

The output of the registration – the transformation can be reapplied to replicate the registered volume. This can be applied in Amira-Avizo or can be independently performed with the transformation is provided in **Table S3.** This transformation is defined following the form of the Class SoTransform from the open inventor toolkit[1] where the geometric 3D transformation consists of (in order) a (possibly) non-uniform scale about an arbitrary point, a rotation about an arbitrary point and axis, and a translation. The transformations can be thought of as being applied in that order, matrices are actually premultiplied in the opposite order. e.g. where the transformation is applied from a combination of the translation matrix (T), the rotation matrix (R) and the scale matrix (S). The affine transformation M=S*R*T is calculated by applying scaling first, then the rotation and finally the translation. Each parameter is described in **Table S2.**

| Parameter name | Description | Default value |
|---|---|---|
| **translation** | Translation vector<br>Field containing a three-dimensional vector. | 0 0 0 |
| **rotation** | Rotation specification<br>4 values represent an axis of rotation followed by the amount of right-handed rotation about that axis, in degrees. | 0 0 1 0 |
| **scaleFactor** | Scale factors<br>Field containing a three-dimensional vector. | 1 1 1 |
| **scaleOrientation** | Rotational orientation for scale<br>Four floating point values separated by whitespace. The 4 values represent an axis of rotation followed by the amount of right-handed rotation about that axis, in degrees. | 0 0 1 0 |
| **center** | Origin for scale and rotation<br>Field containing a three-dimensional vector. | 0 0 0 |

**Table S2. parameter descriptions to define the registered volume**

[1] https://www.openinventor.com/reference-manuals/NewRefMan1030/RefManJava/com/openinventor/inventor/nodes/SoTransform.html

| Dataset | Voxel size | Translation | Rotation | Scale factor | Scale orientation | Centre |
|---|---|---|---|---|---|---|
| **2.6um_LAD AF-2021-17_kidney-right_VOI-03.1** | 5.2 | 6691.73, 24725.4, 28319.2 | 0.000967297 0.00446794, -0.99999 19.9447 | 1 1 1 | 0.00782034, -0.025613, -0.999641 | 5010.2 5010.2 3629.6 |
| **6.5um_LAD AF-2021-17_kidney-right_VOI-03** | 13 | -4068.46, 15889.5, 3561.3 | -0.00163864, -0.0180448, -0.999836, 19.6033 | 1.00354 1.00354 1.00354 | 0.222067, 0.721417, -0.65593 | 12564.5 12207 38701 |

**Table S3. Registration parameters for the high-resolution volumes of interest to the overview whole organ image volume.**

#### 2 Image Processing pipeline

##### 2.1 Validation of segmentation using hierarchical data

Quantification of the manual segmentation accuracy for the overview scans (down sampled to 50 µm/voxel) was provided by using segmentation of the 13 µm/voxel dataset (down sampled from 6.5 µm/voxel ) as a 'ground truth' and using the cl-dice metric [1] to quantify the difference. The cl-DICE metric quantifies the proportions of false positive and false negative vessels in 3D segmentations of vascular networks, by utilising the midline (or skeleton) representation of the segmentation, this reduces the bias for larger vessels to affect the metric score as is the case for a classical DICE metric.

By using the 13 µm/voxel datasets as a 'ground truth' for the 50 µm/voxel dataset (**Figure S3**) it is possible to visually see the overlap between the segmentations for the larger vessels with additional smaller vessels present in the 13 µm/voxel segmentation that are not visible in the 50 µm/voxel. This is quantified by the topological precision (sensitive to false positives) and topological recall, (sensitive to false negatives). The topological precision (related to true positives) was found to be 0.97 and the topological recall (related to false negatives) was 0.5, with the undetected vessels having diameter less than 50 µm, (**Figure S3).** This demonstrates that our approach to manual annotation (utilising three independent annotators in an iterative manner) correctly identifies 97% of vessels $<$50 µm radius across the whole intact human kidney.

##### 2.2 Skeletonization optimisation

Skeletonization reduces a binary image of a vascular network in this case to a 1D skeleton or spatial graph. This can be performed using a number of different algorithms and thus the process should be well described and optimised. We optimised our skeletonisation following the approach of Walsh and Berg [2]. We considered three different skeletonization algorithms: Centerline-Tree (Amira-Avizo), AutoSkeleton (Amira-Avizo), and the Medial Thinning algorithm (implemented in VesselVio [3]).

Firstly, to we choose the most appropriate algorithm we applied the recently defined skeleton super-metric [2] for each of the methods which seeks to minimise the geometric difference between a validated segmentation and the skeleton representation.

The super metric contains 5 components calculated relative to the binary segmentation.

If $I_0$ be the binary image segmentation, we were able to define 5 features

1. Total network volume $V$
2. Euler number of the largest connected component $\chi = N - E$

3. Number of connected components $cc$
4. The DICE score for the branching nodes in randomized sub volume of the network. $B$ (range 0-1)
5. Cl-sensitivity score [1]. $Cl$ (range 0-1)

These features were combined into a single function that can used as a cost function for skeleton optimisation. For any skeleton $S_i$ created from a binary image $I_0$ we write this function as

$$M_{S_i} = \sqrt{\left(\frac{V_{I_0} - V_{S_i}}{V_{I_0}}\right)^2 + \left(\frac{CC_{I_0} - CC_{S_i}}{CC_{I_0}}\right)^2 + \left(\frac{\chi_{I_0} - \chi_{S_i}}{\chi_{I_0}}\right)^2 + \left(\frac{B_{I_0} - B_{S_i}}{B_{I_0}}\right)^2 + \left(\frac{Cl_{I_0} - Cl_{S_i}}{Cl_{I_0}}\right)^2}$$

A skeleton with super metric value 0 can be considered optimal and optimsation can be performed formally using e.g. Latin hyperparameter search, or informally with a random search approach from the user.
In our case metric was used in a two phase process; firstly to choose skeletonisation algorithms and secondly to optimize the chosen algorithm.
In the first stage the Centreline Tree was compared to the Vesselvio and Autoskeleton algorithms and was selected on the basis of the low super-metric value and as it is known to enforce tree topology and thus enable Strahler Ordering analysis.
Once selected the two parameters of the Centreline Tree algorithm: slope and zeroVal were optimized following [2] to find the optimal parameter values.
The value for each component of the super-metric as well as the final metric for each of the initial skeletonization algorithms and for the optimized Centerline Tree algorithm is show in Figure S5

#### 2.3 Multi-scale smoothing

The mulitscale nature of HiP-CT makes it a unique imaging technique; however, when performing skeletonization the magnitude of the multiscale features (ranging from large arteries ca. 1000 µm diameter down to arterioles of ca. 70 µm diameter), is problematic for choosing a single set of parameters for skeletonization. To overcome this challenge, we utilised a Strahler ordering system (**Figure 2 Step 3**) to divide the network into larger and smaller calibre vessels, (those above and below Strahler order 5). We the apply an iterative weighted smoothing to all points in Amira-Avizo, where the smoothed location of any point is given by iteratively calculating weighted average of the current the two neighbour points and the position and. Parameter values were found empirically as

0.8, 0.1 and 15 for the neighbour points, current point and iterations respectively. This reduced the tortuosity in the larger vessels (**Figure 2 Step 4**) which is caused by noise on the surface of the segmented large vessels, and which if uncorrected impacts severely on the correction for collapsed radius vessels.

#### 2.4 Correction of radius value for collapsed vessels

A second correction needed specifically for HiP-CT data is the correction of collapsed vessels. Some collapse of larger vessels during the HiP-CT preparation process is unavoidable (**Figure S6** for examples). Whilst techniques such as microfill, could avoid this issue, microfill can lead to radius over estimation and would drastically increase sample preparation complexity. Additionally, microfill would interfere with the dynamic range sensitivity which is central to achieving such high contrast with HiP-CT. Thus, we chose to identify and correct the collapsed vessels in image post processing.

Collapsed vessels could be identified by visual inspection of the data in some cases. See Figure S6 and are easily distinguished in the spatial graph due to the large underestimate of the radius in these segments (caused by the maximum inscribed sphere estimation for radius)

We delineated two distinct cases of collapsed vessel, one where there is a small collapsed portion in an otherwise patent vessel (Figure S6Bi and Bii), second, where the majority of the vessel is collapsed (Figure S6Cii). The reduction in radius in the skeletonised form of the networks can be seen in Figures S6Bii and S6Cii.
The Centreline Tree algorithm estimates radius of the vessels using a maximum inscribed sphere approach. We choose to retain this approach for non-collapsed vessels for its computational speed and instead to identify and correct only the collapsed vessels. Correction for short subsegment collapsed vessels was performed by plotting radius along each segment, subsegments where the radius was below the $5^{th}$ percentile of the data were replaced with the nearest neighbour.

For larger collapsed vessels we used the Strahler ordering scheme to first identify collapsed vessels (**Figure 2 Step 3**). We plotted Strahler order against mean segment radius (**Figure 2 Step 5**). Outliers in these data (those below the $10^{th}$ percentile) were identified, and the binary image data in a bounding box surrounding the segment were automatically retrieved and presented to an annotator (**Figure 2 Step 6**). It was necessary at this point to have a manual intervention stage where the annotator confirms whether the segment requires correction. In this way we were able to increase the bounds for the outlier detection ($10^{th}$ percentile), to ensure we found collapsed vessels but disregarded vessels that were not collapsed.

Once a segment was confirmed as collapsed the pipeline continued automatically. A new estimate of radius for the larger collapsed vessels was performed by extracting planes from the 3D image data through the segment that were orthogonal to the centreline at each 'point' along the segment. For each of these image planes the perimeter of the segment cross-section was found and converted to a radius with the assumption of a circular cross-section (**Figure 2 step 7**). The final step was to remove the within-segment outliers for these corrected segments, using the same nearest neighbour approach as above but for the 5$^{th}$ and 95$^{th}$ centiles, which result from tortuosity in the sub-segments (**Figure 2 Step 8**). Note this is why multiscale smoothing must be performed before radius correction. After correction the new radii were updated and topological analysis of the skeleton performed.

### 3 Glomeruli Segmentation and Strahler Order Estimation

Estimation of the distance (in Strahler order) from the end of the segmented and skeletonised arterial network was facilitated again by the hierarchical imaging. Using the higher resolution HiP-CT regions of interest we were able to segment individual glomeruli and in some cases like these back to the segmentation of the whole vascular network at 50 µm/voxel. Figure S7 show the same sub region of the kidney segmented and skeletonised at 50, 5.2 and 2.6 µm/voxel. At 2.6 µm/voxel it was possible to link four glomeruli back to the whole vascular network and from there show that there were at two additional Strahler orders between the end of the segmented whole vascular network and these glomeruli. Furthermore we are able to extend this estimation, if we consider that all end points in the 5.2 µm/voxel sub data set branch a further two times before terminating in glomeruli we estimate that glomeruli are between two to four Strahler orders from the end of the 50 µm/voxel dataset.

This estimate can be validated independently when we estimate the total number of glomeruli in this kidney (following a similar approach to glomeruli estimation in HiP-CT data from Walsh et al.) we find the total glomeruli to be approx. 400,000. In the current skeletonisation we have 5666 terminal nodes. Based on the branching ratio we have calculated (2.91) we would need 12 Strahler orders to have the approx. 400,000 end nodes ($2.91^{12}$=368,736) thus and addition 3 Strahler order are predicted on the end of the current tree (which has 9 Strahler Orders).

### 4 Segmentation of compartments of human kidney

| **Hyperparameter** | **Value** |
|---|---|
| First layer size | 64 |
| Layers | 5 |
| Patch size | 64 |
| Stride Ratio | 0.5 |
| Batch size | 128 |

| | |
|---|---|
| Epochs set | 100 |
| Epochs completed | 63 |
| Loss Function | Categorical Cross Entropy |
| Optimiser | AdaDelta |
| Augmentation | 4 times |
| Augmentation types | Flip horz, Flip vert, rot, shear, scale (90-110%) |
| Learning rate | 1 |
| ρ | 0.95 |
| Epsilon | 1.00E-07 |
| Validation % | 20% |
| Patience | 20 |

**Table S4. Hyperparameters of the CNN used for segmentation of human kidney anatomical compartments**

[1] J. C. Paetzold *et al.*, “clDice—A novel connectivity-preserving loss function for vessel segmentation,” in *Medical Imaging Meets NeurIPS 2019 Workshop*, 2019.

[2] C. L. Walsh, M. Berg, H. West, N. A. Holroyd, S. Walker-Samuel, and R. J. Shipley, “Reconstructing microvascular network skeletons from 3D images: what is the ground truth?,” *Comput Biol Med*, vol. 171, p. 108140, 2024.

[3] J. R. Bumgarner and R. J. Nelson, “Open-source analysis and visualization of segmented vasculature datasets with VesselVio,” *Cell Reports Methods*, vol. 2, no. 4, p. 100189, 2022, doi: https://doi.org/10.1016/j.crmeth.2022.100189.
