## Supplementary figures and images for "Mapping the blood vasculature in an intact human kidney using hierarchical phase-contrast tomography"

### Final_ Supplemenrary_figure_S6.tif

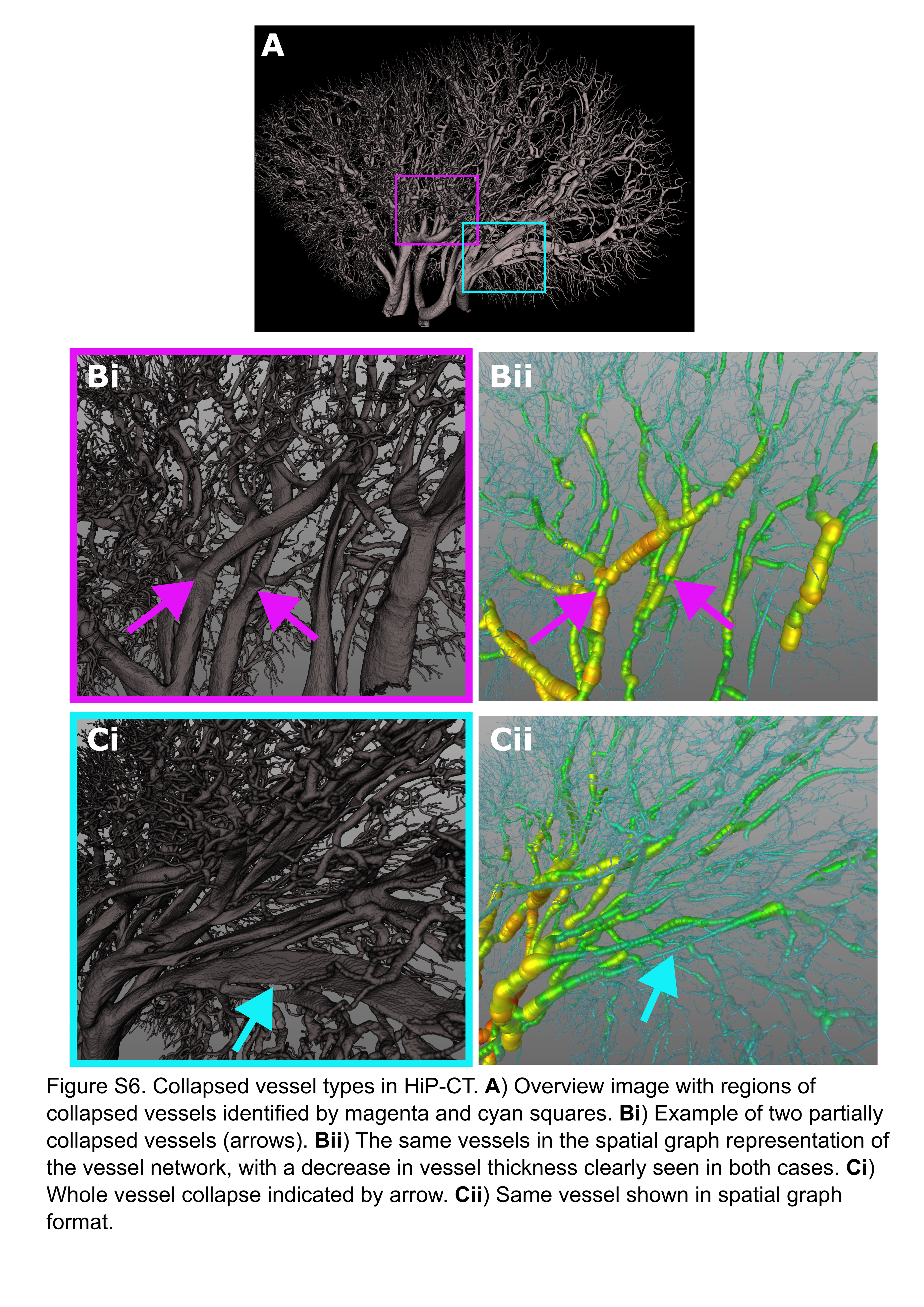

### Final_Supplemenrary_figure_S7.tif

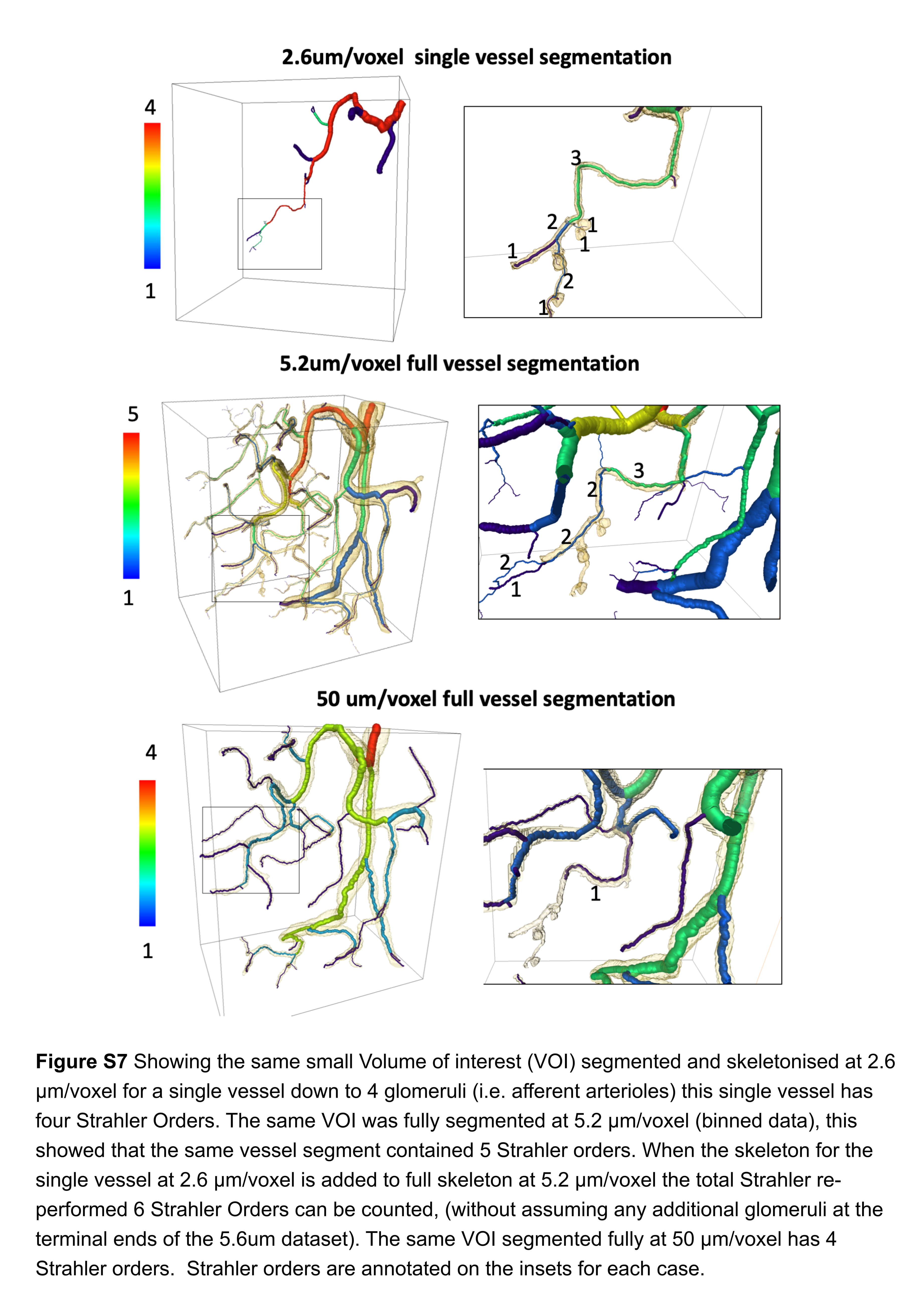

### Final_Supplementary_figure_S1.tif

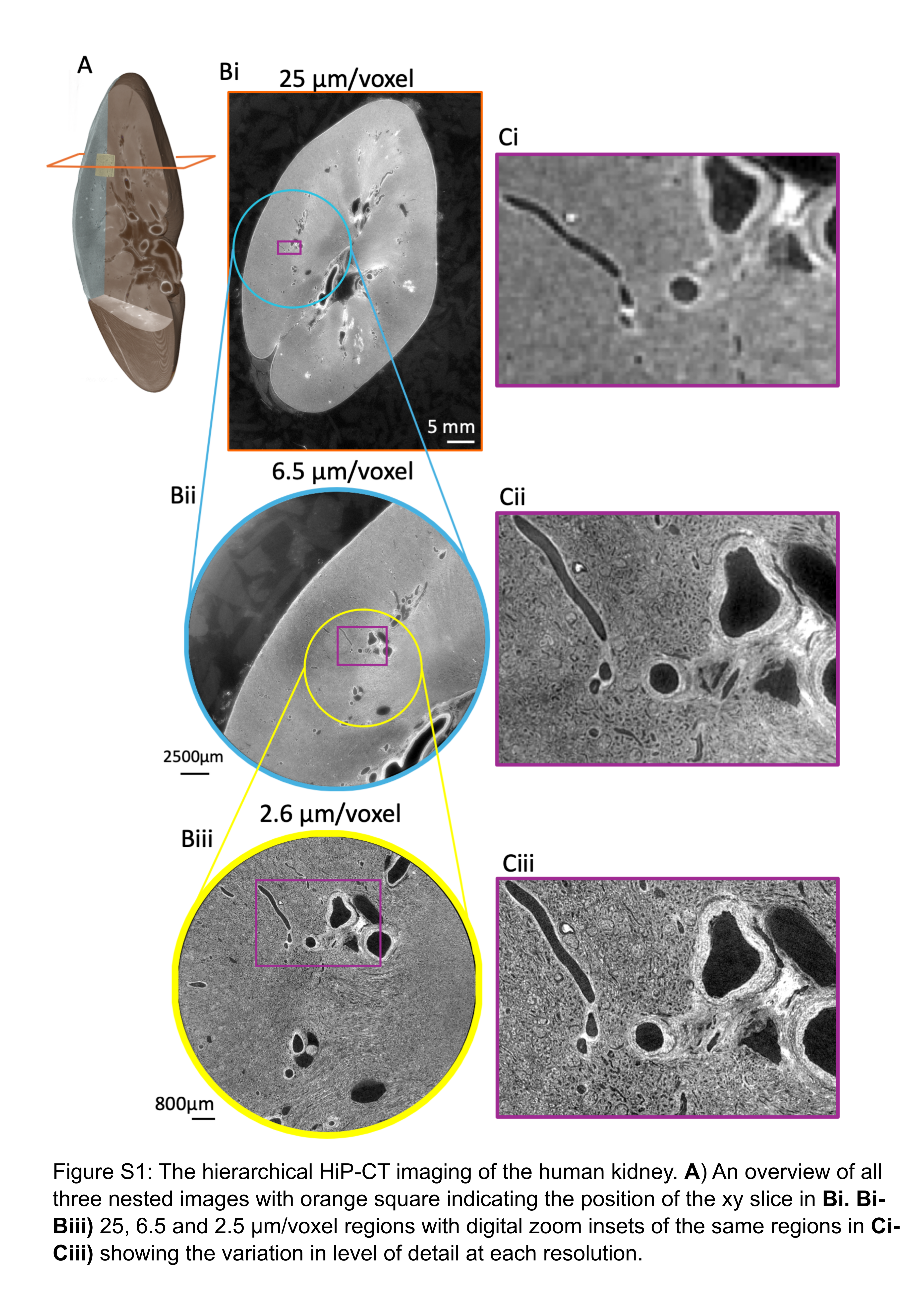

### Final_Supplementary_figure_S2.tif

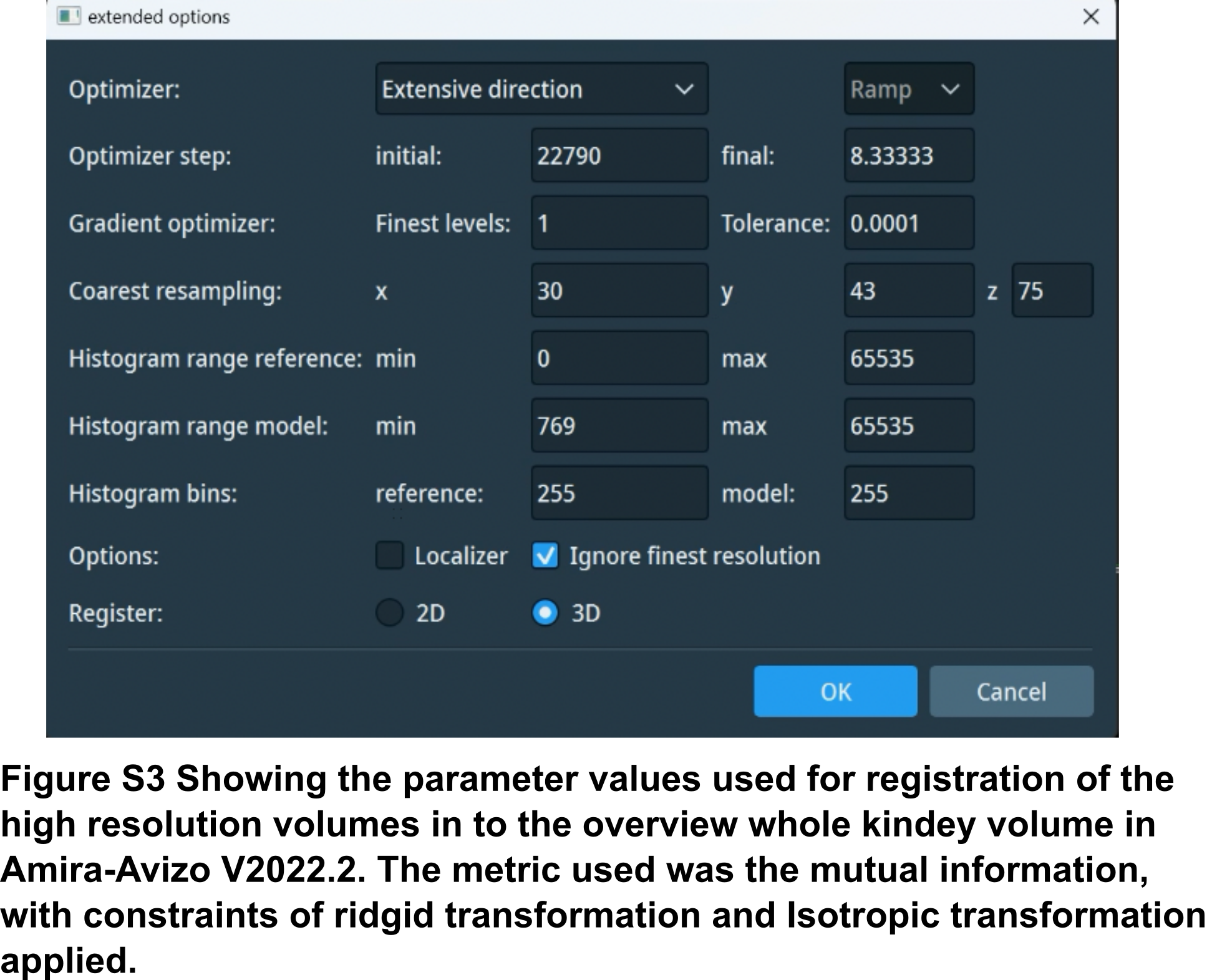

### Final_Supplementary_figure_S3.tif

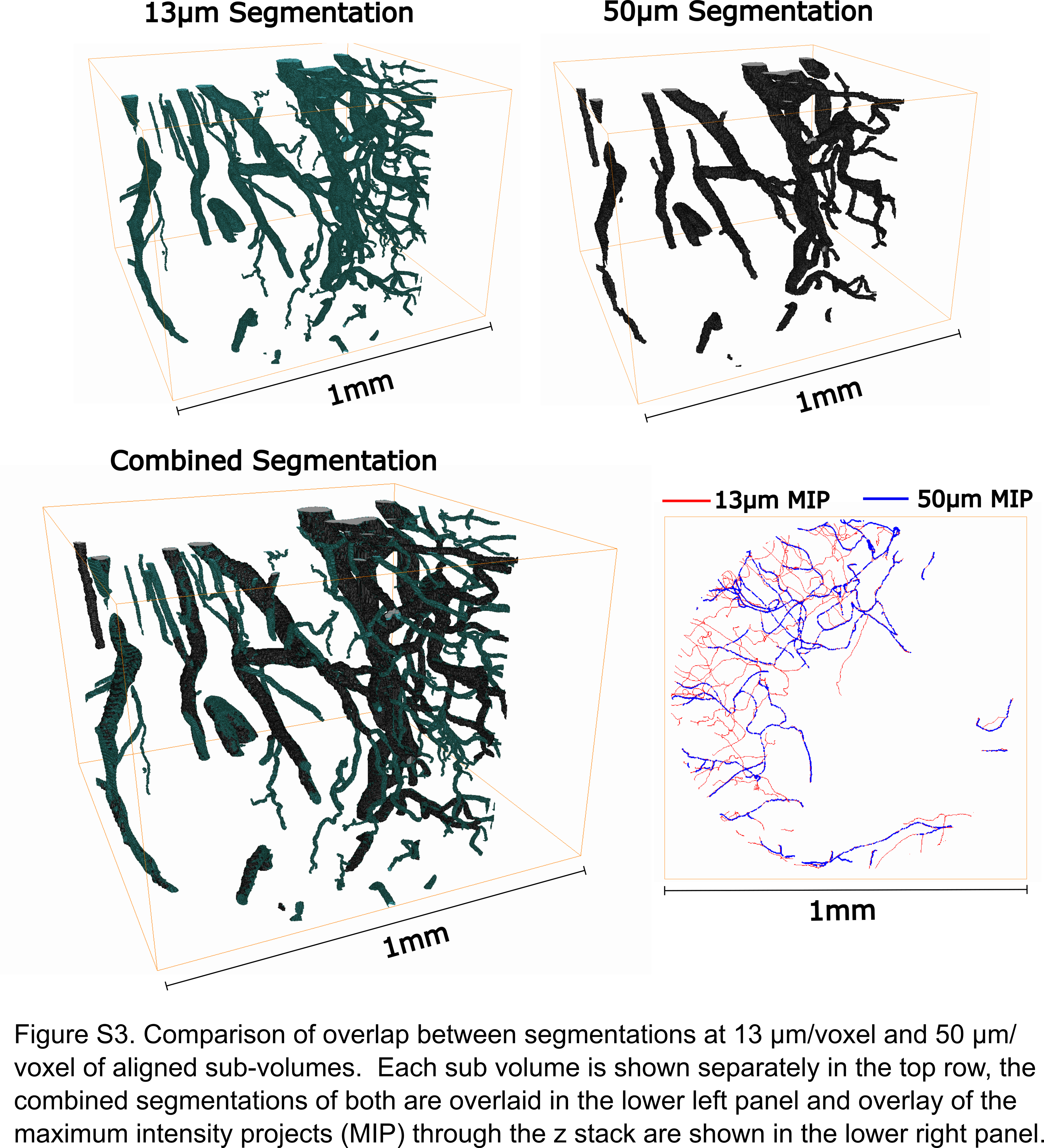

### Final_Supplementary_figure_S4.tif

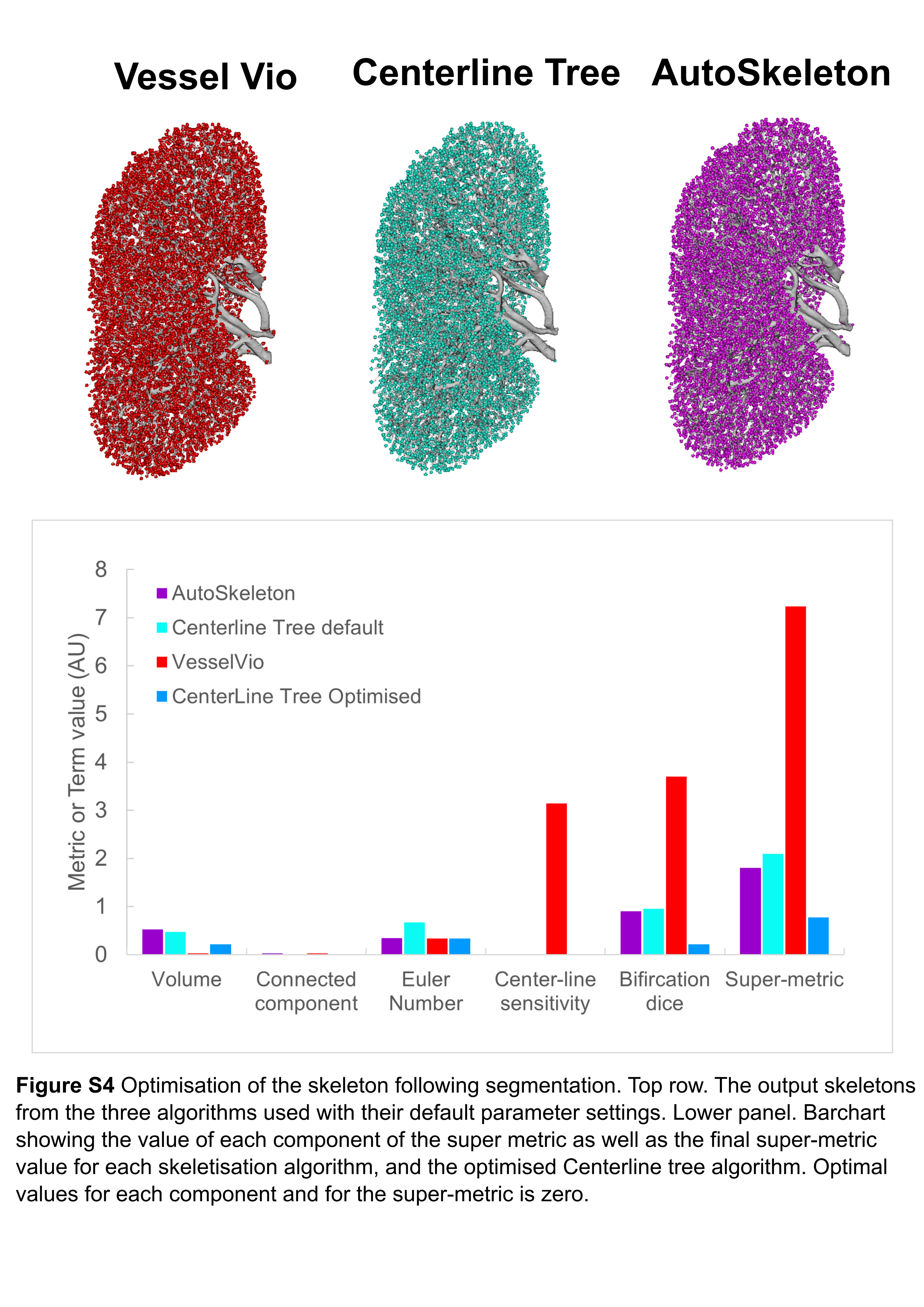

### Final_Supplementary_figure_S5.tif

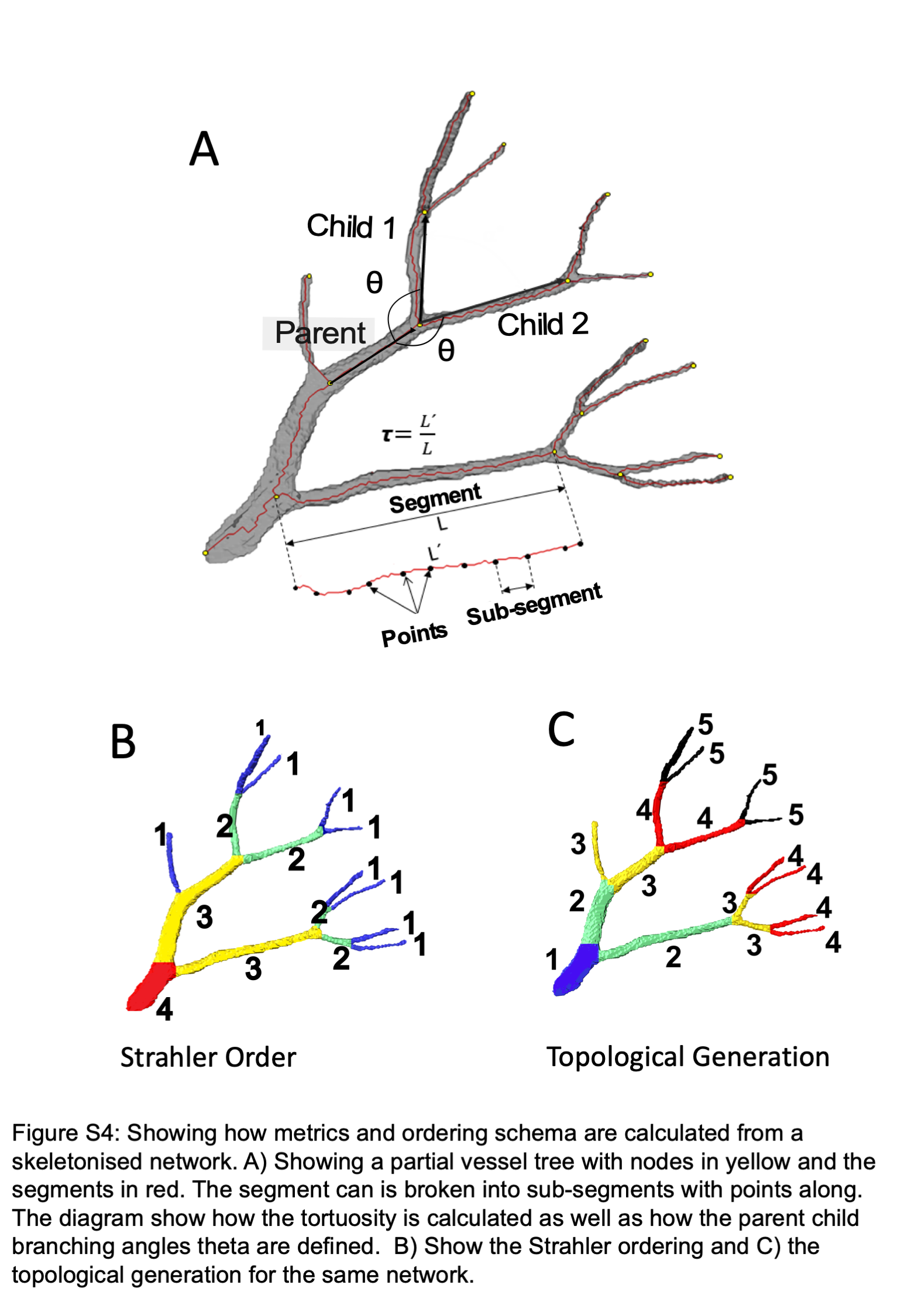
